# The immunometabolic topography of tuberculosis granulomas governs cellular organization and bacterial control

**DOI:** 10.1101/2025.02.18.638923

**Authors:** Erin F. McCaffrey, Alea C. Delmastro, Isobel Fitzhugh, Jolene S. Ranek, Sarah Douglas, Joshua M. Peters, Christine Camacho Fullaway, Marc Bosse, Candace C. Liu, Craig Gillen, Noah F. Greenwald, Sarah Anzick, Craig Martens, Seth Winfree, Yunhao Bai, Cameron Sowers, Mako Goldston, Alex Kong, Potchara Boonrat, Carolyn L. Bigbee, Roopa Venugopalan, Pauline Maiello, Edwin Klein, Mark A. Rodgers, Charles A. Scanga, Philana Ling Lin, Denise Kirschner, Sarah Fortune, Bryan D. Bryson, J. Russell Butler, Joshua T. Mattila, JoAnne L. Flynn, Michael Angelo

## Abstract

Despite being heavily infiltrated by immune cells, tuberculosis (TB) granulomas often subvert the host response to *Mycobacterium tuberculosis* (Mtb) infection and support bacterial persistence. We previously discovered that human TB granulomas are enriched for immunosuppressive factors typically associated with tumor-immune evasion, raising the intriguing possibility that they promote tolerance to infection. In this study, our goal was to identify the prime drivers for establishing this tolerogenic niche and to determine if the magnitude of this response correlates with bacterial persistence. To do this, we conducted a multimodal spatial analysis of 52 granulomas from 16 non-human primates (NHP) who were infected with low dose Mtb for 9-12 weeks. Notably, each granuloma’s bacterial burden was individually quantified allowing us to directly ask how granuloma spatial structure and function relate to infection control. We found that a universal feature of TB granulomas was partitioning of the myeloid core into two distinct metabolic environments, one of which is hypoxic. This hypoxic environment associated with pathologic immune cell states, dysfunctional cellular organization of the granuloma, and a near-complete blockade of lymphocyte infiltration that would be required for a successful host response. The extent of these hypoxia-associated features correlated with worsened bacterial burden. We conclude that hypoxia governs immune cell state and organization within granulomas and is a potent driver of subverted immunity during TB.

## INTRODUCTION

Granulomas are a hallmark of tuberculosis (TB)^1,2^. These spatially organized hubs of immune cells are induced by infection with *Mycobacterium tuberculosis* (Mtb). While granulomas contribute to pathogen containment, they can also support bacterial persistence and immune evasion^3,4^. Whether a granuloma veers towards one extreme or the other hinges on how well the immune system balances tissue-damaging, inflammatory anti-microbial responses against tolerogenic pathways that limit damage but also hinder bacterial clearance. While an optimal balance leads to long-term protection in all granulomas, skewing of this equilibrium can worsen disease even if just a subset of granulomas fails^3–7^.

There is a growing body of evidence to suggest that this immunological balance in granulomas from individuals with active TB disease veers towards tolerance of Mtb bacteria and worsened disease^8–14^. In human TB, we previously demonstrated that macrophages expressing immunosuppressive proteins like IDO1, PD-L1, and TGFβ colocalize with proliferating Tregs in the granuloma myeloid core where Mtb reside^8^. These findings implicate myeloid-mediated immune suppression as a central driver of failed immunity in TB, making it a compelling angle for therapeutic intervention. At present, however, little is known about how these tolerogenic niches are initiated and sustained.

One candidate regulator of immune tolerance in the granuloma is hypoxia, which occurs when local oxygen levels are diminished^15–17^. Immune cell function is inextricably linked to hypoxia via the oxygen-sensitive transcriptional regulator, hypoxia-inducible factor-1 (HIF-1)^18–22^. Because it is constitutively expressed, intracellular levels of HIF-1a (a subunit of HIF-1) are primarily regulated by its rate of degradation, which is highest in normoxia. As oxygen tension drops, HIF-1a stabilization triggers a transcriptional shift towards anaerobic glycolysis^23^. This metabolic shift in cellular metabolism is tightly coupled to dramatic changes in immune function that are generally anti-inflammatory and tolerogenic. For example, in tumors, hypoxia is pathologic and subverts host immunity by suppressing T cell effector functions, inducing of suppressive myeloid cell phenotypes, and promoting preferential infiltration by regulatory T cells (Tregs)^19,24–30^. Although the mechanisms mediating its effects are not well understood, granuloma hypoxia has been associated with reduced drug efficacy, mycobacterial persistence and neutrophil-mediated tissue damage^31–37^.

Hypoxia’s influence on immune regulation in TB granulomas remains unclear, but given the cooccurrence of hypoxic and immunosuppressive signatures in granulomas, we hypothesized that hypoxia imprints a tolerogenic environment that enables Mtb persistence. To address this knowledge gap, we used highly multiplexed imaging (Multiplexed Ion Beam Imaging by Time-of-flight, MIBI-TOF), single-cell RNA sequencing (scRNA-seq), and spatial transcriptomics to interrogate granulomas from Mtb-infected non-human primates (NHP) with known bacterial burdens. Using these techniques, we spatially mapped the immunological and metabolic attributes of these granulomas to identify how hypoxia shapes local cell populations and immune functions that are critical for control of Mtb. We found that the macrophage-rich granuloma myeloid core bifurcates into two metabolic subzones, one of which is hypoxic. This hypoxic environment coincides with macrophages enriched in high bacterial burden granulomas, blunted T cell infiltration, and diminished immune-activating signals. Furthermore, the spatial network of granulomas diverges based on bacterial burden, with bacterially permissive lesions associated with hypoxic cellular niches. Collectively, our data support that the spatial organization of hypoxia-regulated immunometabolic states in a TB granuloma contributes to local control of Mtb persistence and replication.

## RESULTS

### Spatial mapping of NHP TB granulomas with MIBI-TOF

To map TB granuloma composition, spatial organization, and function, we curated a cohort of granulomas from Mtb-infected cynomolgus macaques that included 52 archival specimens from 16 NHPs (**Table S1**). The animals had been infected with a low dose of *Mtb* Erdman (7.4 ± 3.7 CFU) and necropsied between 9-12 weeks post-infection (10.6 ± 1.0 weeks) (**Figure S1A-B**), a timepoint that represents an inflection point of infection where granulomas separate into Mtb-permissive or restrictive lesions^13^. We sampled 1-8 granulomas per animal (3 ± 2 granulomas/animal, **Figure S1C**) where each granuloma had defined characteristics including bacterial burden as measured by colony forming units (CFU), size and inflammatory status as measured by ^18^F-FDG PET/CT imaging, and time of detection post-infection on PET-CT imaging (**Figure S1D**). The 52 granulomas we analyzed spanned a five log_10_-fold range of CFU (0-104400).

From this cohort of tissues, we observed no relationship between granuloma CFU with size, ^18^F-FDG uptake, age, or the histological subtype of granuloma (**Figure S1E-F**). This ensured we could isolate the relationship between granuloma bacterial burden with their cellular and spatial features. Most of the specimens contained necrotic granulomas (caseous) (n = 44) although other histopathological presentations including fibrotic granulomas (n = 14), and non-necrotic granulomas (n = 11) were included in this cohort and often found in the same specimen as multifocal/coalescing lesions (n = 21) (**Figure S1G-H).** We imaged each granuloma with MIBI-TOF (**Figure S2B**), which uses spatially resolved mass spectrometry and metal isotope-labeled antibodies, to obtain 38-plex images of each tissue with subcellular resolution^38^. The antibodies used in these panels were validated for cross reactivity against cynomolgus macaque proteins (**Figure S2A, Table S2**) and the data from these images were extracted, denoised, and normalized using our open-source MIBI-TOF pipeline, Toffy (**Figure S3, Methods**)^39^. The resulting dataset of 1,979 images across 52 specimens and 16 animals presents the most comprehensive spatial profiling of NHP TB granulomas to date and allowed us to explore relationships between cellular composition, spatial patterning, immunotopography, and granuloma-level bacterial control (**Figure 1A**).

**Figure 1:**
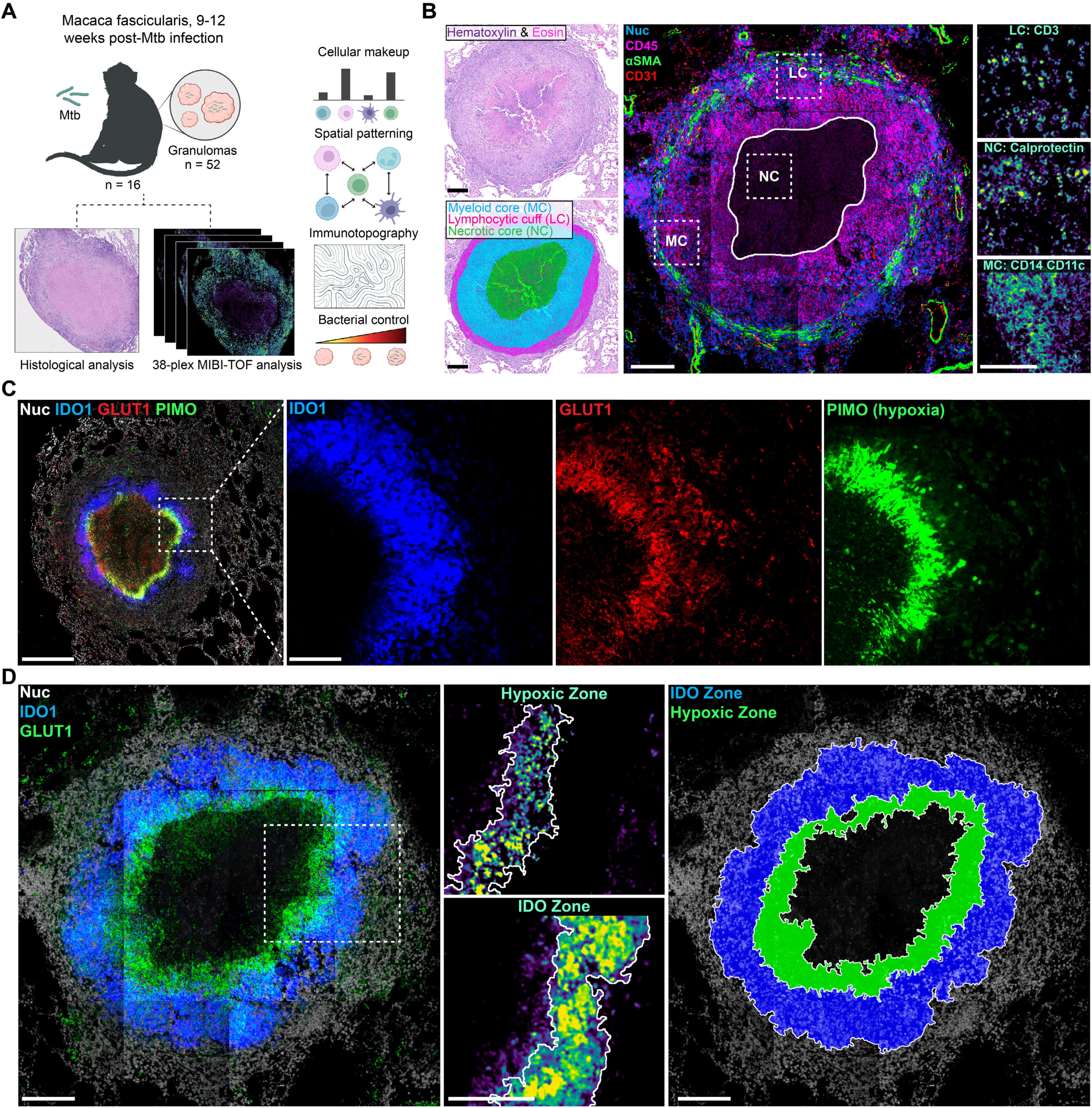
The granuloma myeloid core is comprised of two metabolic subzones. **A)** Conceptual overview of MIBI-TOF analysis of NHP TB granulomas. **B)** Hematoxylin and eosin-stained image of a representative NHP TB granulomas (upper left) with histological zone annotations overlaid (lower left). Center image shows the same granuloma analyzed via MIBI-TOF (blue = nuclei, magenta = CD45, green = αSMA, red = CD31) with white dashed boxes indicating histological subzones (MC = myeloid core, NC = necrotic core, LC = lymphocytic cuff) shown in the adjacent insets (right). All scale bars represent 200 μm. **C)** A NHP TB granuloma analyzed via immunofluorescence microscopy (white = DAPI, blue = IDO1, red = GLUT1, green = PIMO). Scale bars represent 500 μm (large image) and 100 μm (insets). **D)** MIBI-TOF image granuloma in 1A showing IDO1 (blue) and GLUT1 (green) expression (white = nuclei, left) with white dashed box indicating insets with metabolic zone signal (middle), and masked zones overlaid (blue = IDO zone, green = hypoxic zone, right). Scale bars represent 200 μm.

#### The granuloma myeloid core comprises two metabolic subzones

Necrotic TB granulomas are typically described as being comprised of three concentric histological zones^1,40^. The central necrotic core contains debris from dead macrophages and neutrophils alongside Mtb and is surrounded by a macrophage-dominated region referred to here as the myeloid core. These two regions are surrounded by an outermost ring of lymphocytes and other cell types known as the lymphocytic cuff (**Figure 1B**).

It has been previously reported that TB granulomas can become hypoxic, particularly when granulomas undergo caseation^17^. To determine whether granuloma hypoxia is restricted to one of the three canonical spatial compartments in a granuloma, we imaged samples from animals that were administered the *in-situ* hypoxia sensor, pimonidazole (PIMO, **Figure 1C**). We found hypoxic cells were restricted to a sharply demarcated concentric ring that tightly surrounded the necrotic center. Consistent with this, these cells co-expressed GLUT1, a glucose transporter upregulated during hypoxic metabolism (**Figure 1C**)^41^. In previous work, we and others found a similar region of the myeloid core in TB granulomas strongly expressed IDO1—an enzyme that exerts its potent immunosuppressive effects by converting tryptophan into kynurenine^8,42,43^. On examining the spatial distribution of IDO1, PIMO, and GLUT1 co-expression, we find that hypoxia and IDO1 demarcate two distinct subzones of the myeloid core: the IDO zone and the hypoxic zone (**Figure 1C-D**). The IDO zone is positioned in the outer region of the myeloid core, while the hypoxic zone comprises the innermost ring of peri-necrotic macrophages. GLUT1 and PIMO are co-expressed with IDO1 at the transitional interface of these two zones before IDO1 expression sharply drops off in the hypoxic zone (**Figure 1C-D**). In our MIBI-TOF dataset we used GLUT1 and IDO1 expression to systematically annotate these two zones and found that the IDO zone is present in all granulomas (**Figure 1D, Figure S4A-C**). The hypoxic zone is only present in granulomas with necrosis (n = 44 / 52) and as necrosis increases, the hypoxic zone makes up a greater portion of the myeloid core (**Figure S4D**). Based on these observations, we revise the histological zones of the granuloma to reflect this clear and universal metabolic sub-zonation of the myeloid core as: the necrotic core, the hypoxic zone, the IDO zone, and the lymphocytic cuff.

#### Granuloma cell composition diverges across metabolic zones and associates with bacterial control

Using this framework for granuloma microanatomy, we sought to determine how cellular composition and function relates to granuloma metabolic zonation and control of bacterial burden (**Figure 2A**). To do this we used previously validated spatial analytics to map granuloma composition at the pixel, cellular, and multicellular length scales. First, we used the deep learning-based algorithm, Mesmer, to segment each granuloma into single cells and then manually annotated the multinucleated giant cells (**Figure S5A-B**)^44^. We then used Pixie to cluster the pixels in each image to quantify the expression of phenotypic markers for cell lineages (**Figure S5C**)^45^. The pixel clusters and segmented cells from necrotic areas of the granuloma were filtered out to focus our analysis on live cells (**Figure S5D**). Lastly, the segmented cells were clustered into one of 19 cell subsets based on their pixel cluster makeup (**Figure 2B, Figure S5E**). Cell identities were overlaid back onto the segmentation masks to generate cell phenotype maps (CPMs) linking each cell back to its spatial positioning in the granuloma. In total we enumerated 1,202,041 single cells (23,166 ± 12,362 cells per granuloma), assigning 96.9% of them to one of 19 cell subsets ranging in frequency from <1% to 11.9% in aggregate (**Figure 2B, Figure S6A-C**).

**Figure 2:**
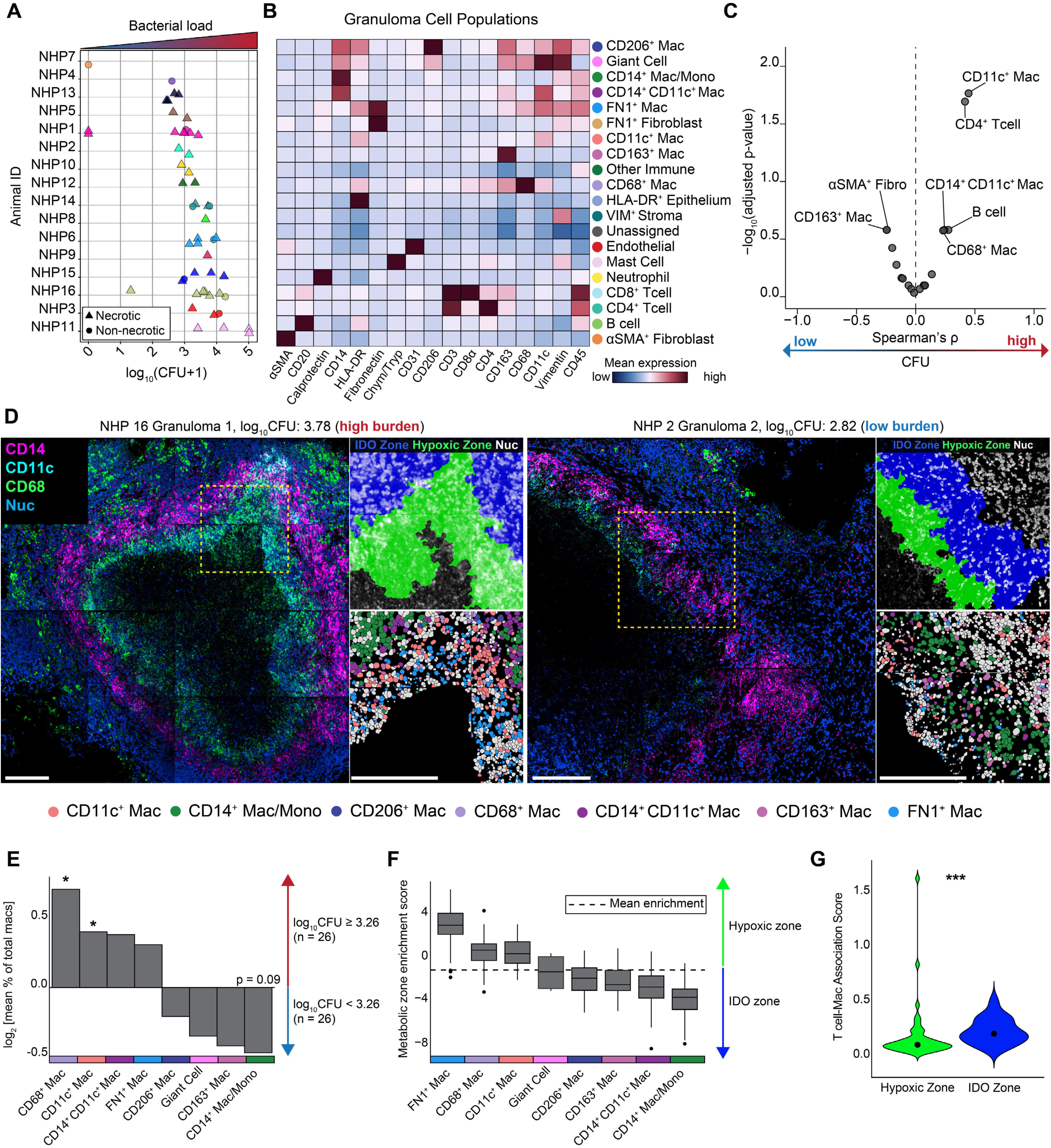
Analysis of granuloma cell composition with respect to metabolic zonation and bacterial burden. **A)** Bacterial burden (log_10_[CFU+1]) per granuloma grouped by NHP. Each color represents a unique animal, and animals are ordered by ascending mean CFU. **B)** Cell phenotype assignments (rows) with normalized expression of phenotypic markers (heatmap columns, z-scored). Rows and columns are hierarchically clustered (Euclidean distance, complete linkage). **C)** Volcano plot of correlation between cellular frequency and granuloma CFU. Plot shows the Spearman rho by FDR-adjusted (5%) p-values. **D)** Representative images of a high (left) and low (right) bacterial burden granulomas. Main image shows macrophage marker expression (blue = nuclei, cyan = CD11c, green = CD68, magenta = CD14). Yellow dashed box delineates insets shown to the right with the metabolic zones (top, green = hypoxic, blue = IDO) and macrophage cell phenotype map (grey = non-macrophage, bottom). Scale bars represent 200 μm. **E)** Log fold-change of mean macrophage frequency (of total macrophages) between granulomas split by CFU threshold (3.26). **F)** Metabolic enrichment score across all macrophage subsets. Dashed line indicates the mean enrichment for all macrophages. **G)** T cell-macrophage interaction score in the hypoxic (green) and IDO (blue) zones. *P* values were calculated with a Wilcoxon rank-sum test (two tailed) (*P < 0.05; **P < 0.01; ***P < 0.001).

Based on the cell clusters enumerated by this workflow, we next compared granuloma composition with CFU to determine which phenotypes associated with bacterially permissive versus restrictive granulomas. We observed that the frequency of two key cell populations was significantly correlated with higher bacterial numbers: CD4^+^ T cells (Spearman rho = 0.42, adj. p = 0.02) and CD11c^+^ macrophages (CD11c^+^ Macs), Spearman rho = 0.45, adj. p = 0.02) (**Figure 2C**). We also found that the frequencies of CD14^+^CD11c^+^ Macs, CD68^+^ Macs, and B cells increased with CFU per granuloma while the frequencies of CD163^+^ Macs and αSMA^+^ Fibroblasts increased with lower CFU, but these trends did not reach statistical significance (**Figure S6D**).

Given that a macrophage subset exhibited the strongest correlation with bacterial burden we more deeply dissected the composition and spatial distribution of each granuloma’s macrophage compartment to identify signatures of permissive versus controlling lesions **(Figure 2D-F**). We found CD68^+^ Macs and CD11c^+^ Macs were elevated in high bacterial burden granulomas (log_10_CFU ≥ 3.26), while less differentiated macrophages (CD14^+^ Mac/Monos) were modestly increased in low burden granulomas (log_10_CFU < 3.26) (p = 0.05, **Figure 2E, Figure S6E**). We next asked whether these macrophage subsets were preferentially enriched in one of the metabolic subzones of the myeloid core. We found a striking bifurcation where high bacterial burden-associated macrophages, such as CD11c^+^ Macs and CD68^+^ Macs were enriched in the hypoxic zone, while CD14^+^ Mac/Monos were the most strongly enriched in the IDO zone (**Figure 2F, Figure S7B**). CD14^+^CD11c^+^ Macs also displayed a spatial preference for the IDO zone, while a population of fibroblast-like macrophages (FN1^+^ Macs) were stringently restricted to the hypoxic zone (**Figure 2F, Figure S7A-C**).

We next performed a similar analysis to understand how these metabolic zones relate to T cell infiltration by enumerating a T cell-macrophage spatial association score across metabolic zones. While T cells mainly localized to the lymphocytic cuff (**Figure S7A**), those that infiltrated the myeloid core were primarily present in the IDO zone, which is positioned most closely to the lymphocytic cuff. T cell infiltration into the hypoxic zone was substantially lower, particularly for CD8^+^ T cells and we observed few T cell-macrophage associations in this region (**Figure 2G, Figure S7D**). Collectively, these data indicate that the hypoxic zone is enriched for macrophages associated with elevated bacterial loads and limited T cell infiltration suggesting that hypoxic conditions in this region contributes to Mtb-permissive rather than Mtb-restrictive granulomas.

#### Macrophage metabolism is coupled to spatial positioning and functional state

Having identified a macrophage signature linking metabolic microenvironment and bacterial control in granulomas, we sought to determine how metabolism affects macrophage phenotype, position, and function. To do this we turned to a single-cell RNA sequencing dataset (scRNA-seq) where granulomas were sampled from cynomolgus macaques after 10 weeks of Mtb infection, a similar timeframe as the samples that we imaged with MIBI-TOF^46^. To identify the macrophage populations that are most likely to be present in the IDO and hypoxic zones, we evaluated expression of two complementary glucose transporters *SLC2A1* (GLUT1) and *SLC2A3* (GLUT3), and *IDO1* in the six original macrophage clusters from this dataset. *SLC2A3/GLUT3* expression was included to compensate for the low abundance of *GLUT1* transcript in this dataset. To bolster our classification criteria, we then assessed the metabolic pathway score for glycolysis in these clusters (**Figure 3A-B**). Much like our MIBI-TOF dataset, *IDO1* expression was elevated across two clusters (**Figure S7E**), but we found that just one of these clusters expressed a gene program consistent with hypoxia and glycolysis. We annotated these as IDO zone macrophages (macs) and hypoxic zone macs, respectively.

**Figure 3:**
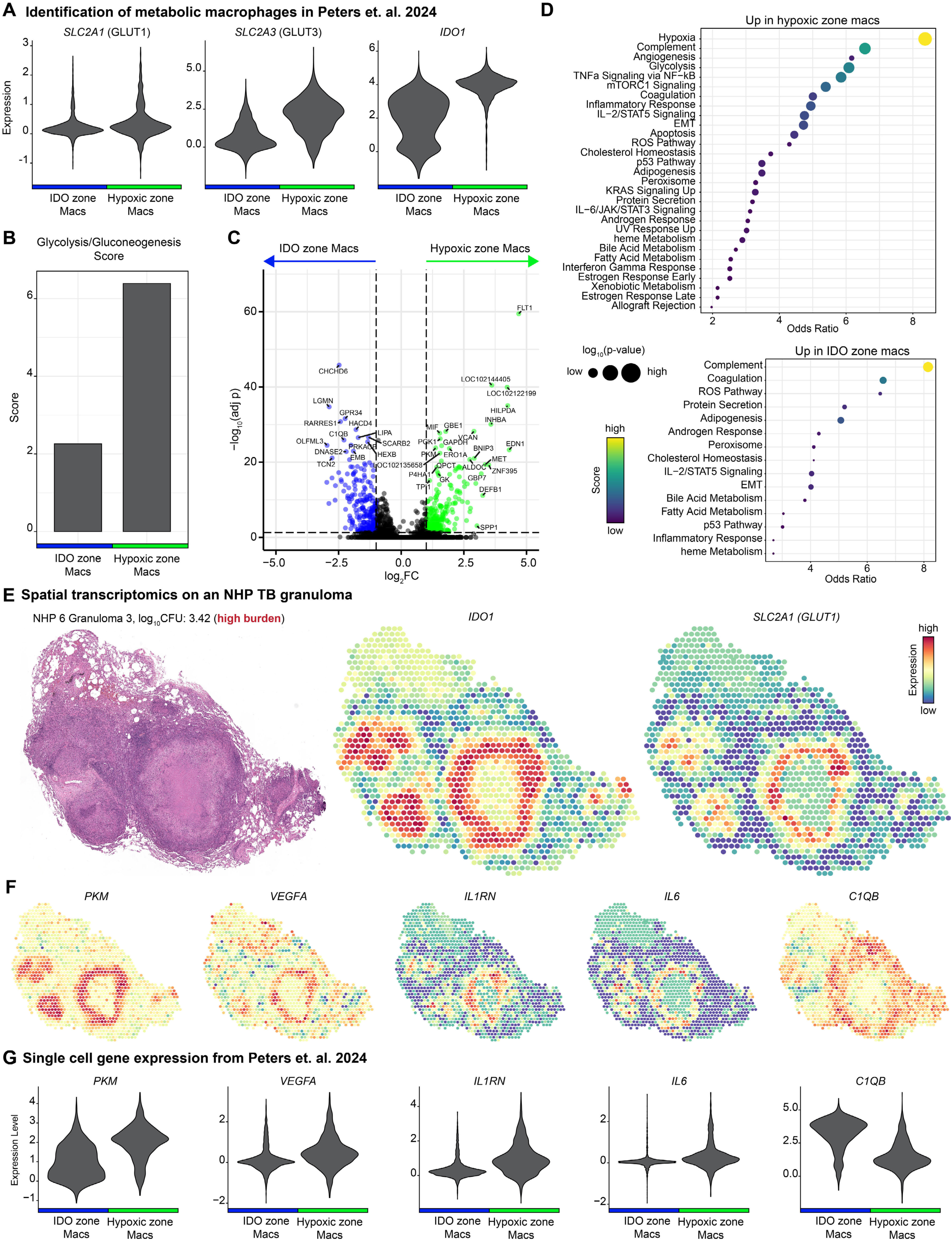
Transcriptomic analysis of cellular metabolism and functional state of granuloma macrophages. **A)** Expression of *SLC2A1*, *SLC2A3*, and *IDO1* in IDO zone (blue) and hypoxic zone macrophages (green). **B)** Metabolic score for glycolysis/gluconeogenesis. **C)** Volcano plot showing differential gene expression for genes upregulated in hypoxic zone (green, right) versus IDO zone (blue, left) macrophages. Colored dots indicate genes with log2 fold-change ≥ 0.5 and adjusted p-value < 0.05. **D)** Gene set enrichment analysis of pathways upregulated in hypoxic zone (top) and IDO zone (bottom) macrophages. Pathways are ordered by descending odds ratio. Dot size is proportional to significance and color is proportional to the pathway score. **E)** Hematoxylin & eosin staining of an NHP TB granuloma analyzed with Visium. Expression of *IDO1* and *SLC2A1* overlaid onto the analyzed spatial regions (spot diameter = 50 μm**). F)** Expression of key genes overlaid onto spots analyzed via Visium. **G)** Expression of genes shown in F in IDO zone versus hypoxic zone macrophages.

To gauge the relationship between the metabolic and functional states of these distinct macrophage groups, we performed differential gene expression analysis and gene set enrichment analysis (**Figure 3C-D**). To spatially validate the results of this approach, we also performed a 10x Visium spatial transcriptomic analysis on a granuloma from our MIBI-TOF dataset (**Figure 3E-F**). We found that hypoxic zone macs upregulated functional programs with glycolysis, hypoxic signaling, and angiogenesis, while IDO zone macs were characterized by upregulation of the complement pathway and ROS production (**Figure 3D**). Our Visium dataset validated the concentric organization of the IDO and hypoxic zones of the myeloid core (**Figure 3E**). Furthermore, in line with our scRNA-seq analysis we observed that, relative to IDO zone macs, hypoxic zone macs expressed higher levels of pyruvate kinase (*PKM*) and vascular endothelial growth factor (*VEGFA*) consistent with a hypoxic state, along with *IL1RN*, a driver of immune suppression, and *IL6* which is associated with worsened TB disease^11,46^. IDO zone macs, on the other hand, showed upregulation of complement pathway factor *C1QB*, which extended into the lymphocytic cuff. The divergent transcriptional profiles of these two macrophage populations— despite their close spatial proximity—underscores the role metabolism may be playing in imprinting macrophage function in granulomas with hypoxia-favoring immunosuppressive macrophage signatures.

#### Radial immunotopography analysis links cellular and functional granuloma microenvironments

Having observed that metabolism underlies macrophage function and organization, we next asked how metabolic microenvironments influence the global organization of functional cell states in a granuloma. To do this, we turned to the field of Geographic Information Systems (GIS) and adapted a geospatial technique known as buffer analysis^47,48^. In this approach, a series of nested, concentric rings (buffers) radiating out from user-specified landmark are first generated via geodesic expansion. Then, the local composition within each of these buffers is compared to understand how the radial distribution of features is coordinated with respect to distance from the landmark. Here, we defined landmarks as the center of the granuloma (either necrotic core or centroid) and generated buffers around these centers (**Figure 4A**).

**Figure 4:**
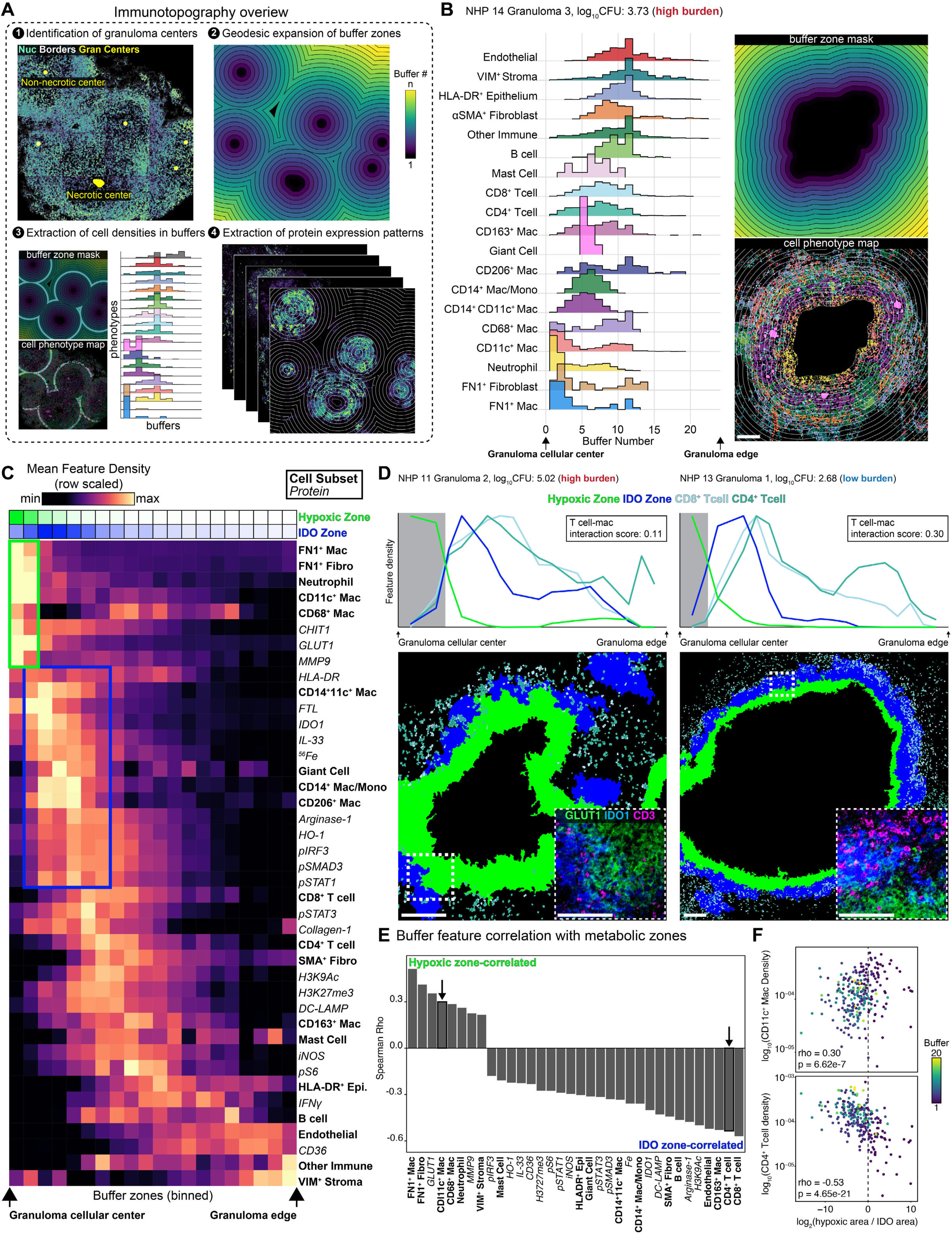
Topographical analysis of granuloma radial structure. **A)** Conceptual overview of immunotopography of NHP TB granulomas. **B)** Cellular topography analysis for a representative granuloma. Histograms represent the density of each cell subset in each buffer zone. Cells are ordered by relative density from the granuloma center (bottom-left) to the outer edge of the granuloma (top-right). Images on the right sown the buffer mask and buffer zones overlaid on the cell phenotype map. Scale bar represents 200 μm. **C)** Heatmap of feature density (rows) in binned buffer zones (columns) for all granulomas. Value represents the mean density per feature and rows are ordered by max density across each buffer zone. Green and blue boxes indicate buffer zones associated with the hypoxic and IDO zone, respectively. **D)** Track plots (top) and representative images (bottom) of the radial distribution of CD4+ T cells (teal), CD8+ T cells (light blue), the hypoxic zone (green), and the IDO zone (blue) across buffer zones. Track plots display the T cell-macrophage interaction score for each granuloma displayed. Gray box indicates switch from hypoxic to IDO zone. Zoomed insets in images display expression of GLUT1 (green), IDO1 (blue), and CD3 (magenta). Scale bars represent 200 μm. **E)** Statistically significant correlations between cellular and protein features and metabolic enrichment across buffers. Features are ordered by descending Spearman rho value**. F)** Plots displaying boxed features from E versus metabolic enrichment score. Y-axes are log-scaled, dots are colored by buffer number. *P* values were calculated with a Student’s t-Test.

Using this approach, termed here as immunotopography, we evaluated the density of cell subsets and the expression of functional markers across the buffers of each granuloma (**Figure 4B, Figure S8A-B**). We then created a consensus map of granuloma radial topography for understanding how both protein expression and cell densities are topographically distributed across the granuloma’s spatial landscape (**Figure 4C**). This global topographic summary revealed a well-defined spatial encoding of immunometabolic traits of a granuloma. The innermost topographic module of the granuloma is hypoxic zone, which is associated with high neutrophil density and expression of MMP9 and CHIT1 in addition to macrophages enriched in high CFU granulomas and GLUT1 (**Figure 4C**). Further supporting an immunosuppressive program of hypoxic zone macrophages, we see that GLUT1 expression is associated with downregulation of HLA-DR (**Figure 4C, Figure S8C**).

As a granuloma transitions to the IDO zone, we observed upregulation of heme metabolism (^56^Fe and FTL) as well as IL-33 production. We also found that the IDO zone is depleted of the histone modification, H3K9Ac (**Figure 4C, Figure S8D**). This unique epigenetic state may explain, in part, how metabolism imparts a distinct functional state on IDO zone macs despite their close spatial proximity to the hypoxic zone and lymphocytic cuff. Along these lines, the IDO zone shows a strong skewing toward pSTAT1, suggestive of ongoing type 1 interferon signaling, while the hypoxic zone displayed a pIRF3 and pSTAT3 dominant signature of cytokine signaling (**Figure S8E**). We also found that the IDO zone is subdivided into adjacent pockets of IDO1-expressing cells encircled in collagen-1 (**Figure 4C, Figure S8F**) suggesting this physical encapsulation promotes the IDO zone’s unique attributes. Lastly, signatures of T cell activation and inflammation such as pS6, iNOS, IFNγ, and DC-LAMP appeared spatially segregated from the innermost region of the granuloma myeloid core, indicating that these critical immune responses are segregated from the primary spatial niche where macrophages are most likely to be interacting with bacteria (**Figure 4C**).

Using immunotopography we can clearly see the halted infiltration of T cells into the hypoxic zone (**Figure 4D**). To further quantify radial relationships of granuloma cellular and spatial features, including lymphocytes, with the metabolic switch from the IDO zone to the hypoxic zone we conducted buffer-based spatial correlation analysis (**Figure S9**). This revealed that numerous features correlated with the metabolic switch from the IDO zone to the hypoxic zone of the granulomas, most notably T cells that are spatially sequestered away from areas of hypoxia (**Figure 4E-F**). This emphasizes the impact that local metabolic activity has on global organization and function of the granuloma.

#### Spatial cellular network configuration underlies controlling versus permissive TB granulomas

Topographical analysis of the granulomas uncovered spatial patterns suggesting that the radial organization of a granuloma subverts the necessary immune responses for bacterial control, especially in the presence of hypoxia. However, this cohort of granulomas represents a large range of CFU, indicating that it comprises granulomas that have controlled infection, those that are permissive for Mtb, and some that may be on the trajectory of control or loss of control at the time of collection. To identify the cellular interactions that distinguished granulomas controlling infection from those permitting bacterial persistence we leveraged QUantitative InterCellular nicHe Enrichment (QUICHE), an algorithm that identifies cellular interactions enriched in controlling versus permissive granulomas (**Figure 5A-B**)^49^. This approach unveiled 98 differentially abundant cellular niches in high burden (log_10_CFU ≥ 3.26) versus low burden (log_10_CFU < 3.26) granulomas (**Figure S10A**). From these niches we generated spatially weighted networks of the interactions defining these two groups of granulomas (**Figure 5A**).

**Figure 5:**
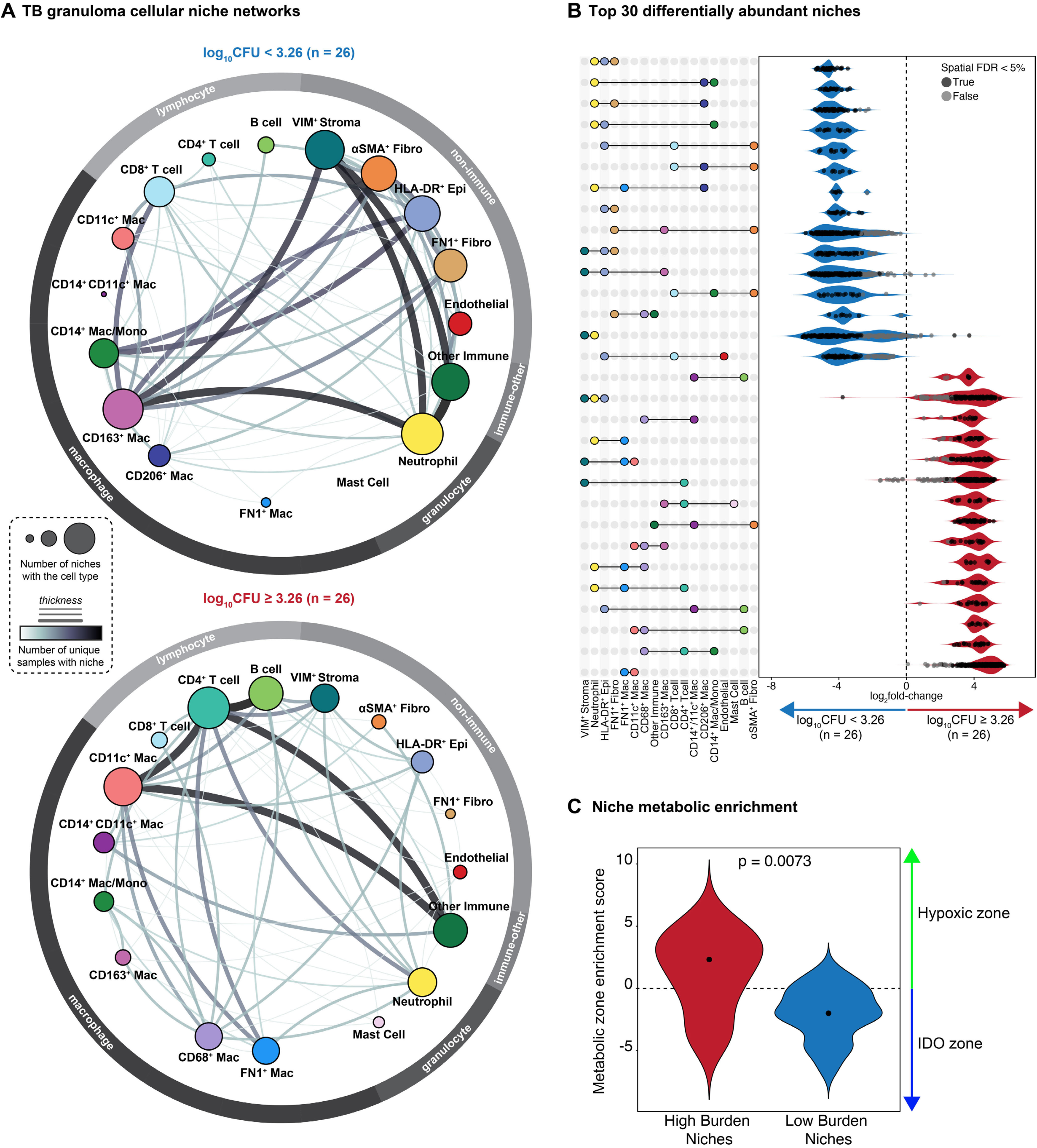
Spatial analysis of cellular niches and networks in high versus low bacterial burden TB granulomas. **A)** Cellular networks derived from differentially abundant niches. Each node represents a cell type. Node size is proportional to connectedness, as measured by eigenvector centrality. Edge thickness is proportional to the number of unique samples with the corresponding interaction**. B)** Top 30 differentially abundant cellular niches in high versus low bacterial burden (based on log_10_CFU threshold 3.26). Shading of dots indicates significance derived from spatial FDR (5%). **C)** Metabolic enrichment score of spatial niches broken down by bacterial burden.

By comparing these two networks we identified several themes of granuloma cellular connectivity and bacterial control. First, we found that low bacterial burden granulomas had more tightly connected spatial niches dominated by CD8^+^ T cells, fibroblasts, CD14^+^ Mac/Monos, CD163^+^ Macs, and CD206^+^ Macs. On the other hand, high burden granuloma niches were dominated by hypoxic zone-associated CD11c^+^ Macs and CD68^+^ Macs. Secondly, we found that endothelial cells were more abundant in low burden granuloma niches despite the frequency of endothelial cells having no correlation with granuloma CFU (**Figure 5A, Figure S6D**). This suggests that increased vascularization within the immune infiltrated region of the granuloma may mitigate the effects of hypoxia and its immunosuppressive consequences. Lastly, we observe that neutrophils appear in niches of both high and low burden granulomas. However, these neutrophils appear in two distinct locations of granulomas. High burden granulomas are enriched for neutrophils in the center of the granuloma, like those associated with the hypoxic zone, while low burden granulomas have increased neutrophil presence in the outer regions of the granuloma (**Figure 5A, Figure S10B**). In line with this, we observe that the level of neutrophil-associated protein calprotectin is higher in the necrotic core of high bacterial burden granulomas (**Figure S10C)**.

Cells also displayed niche-specific functional states (**Figure S10D**). For example, we found that that CD11c^+^ Macs and CD68^+^ Macs in low burden niches upregulated expression of Arginase-1, while these two populations favored expression of GLUT1 in high burden niches (**Figure S10D**). This paired with the cellular composition of the niches point to hypoxia’s outsized influence on cellular connectivity in high bacterial burden granulomas. To test this, we evaluated the spatial localization of each niche across the IDO and hypoxic granuloma zones and saw that the spatial niches abundant in high CFU granulomas displayed a clear enrichment in the hypoxic zone of the granuloma (**Figure 5C**). By enumerating cellular niches differentially abundant in the IDO zone versus hypoxic zone of all granulomas (**Figure S11A**), we observed that the cellular network of high bacterial burden granulomas most resembled that of the hypoxic zone.

Overall, we find that the granuloma’s spatial design is tightly coupled with immunometabolic zonation. By evaluating archival tissue of a pulmonary TB specimen from a human patient, we confirm that the same metabolic zonation of the myeloid core and associated spatial patterning of lymphocytes and macrophages is present in human disease (**Figure 6A**). Our findings indicate that hypoxia imprints an environment in NHP and human TB granulomas that is permissive to bacterial persistence, stalled T cell infiltration, and diminished immune activation (**Figure 6B**).

**Figure 6:**
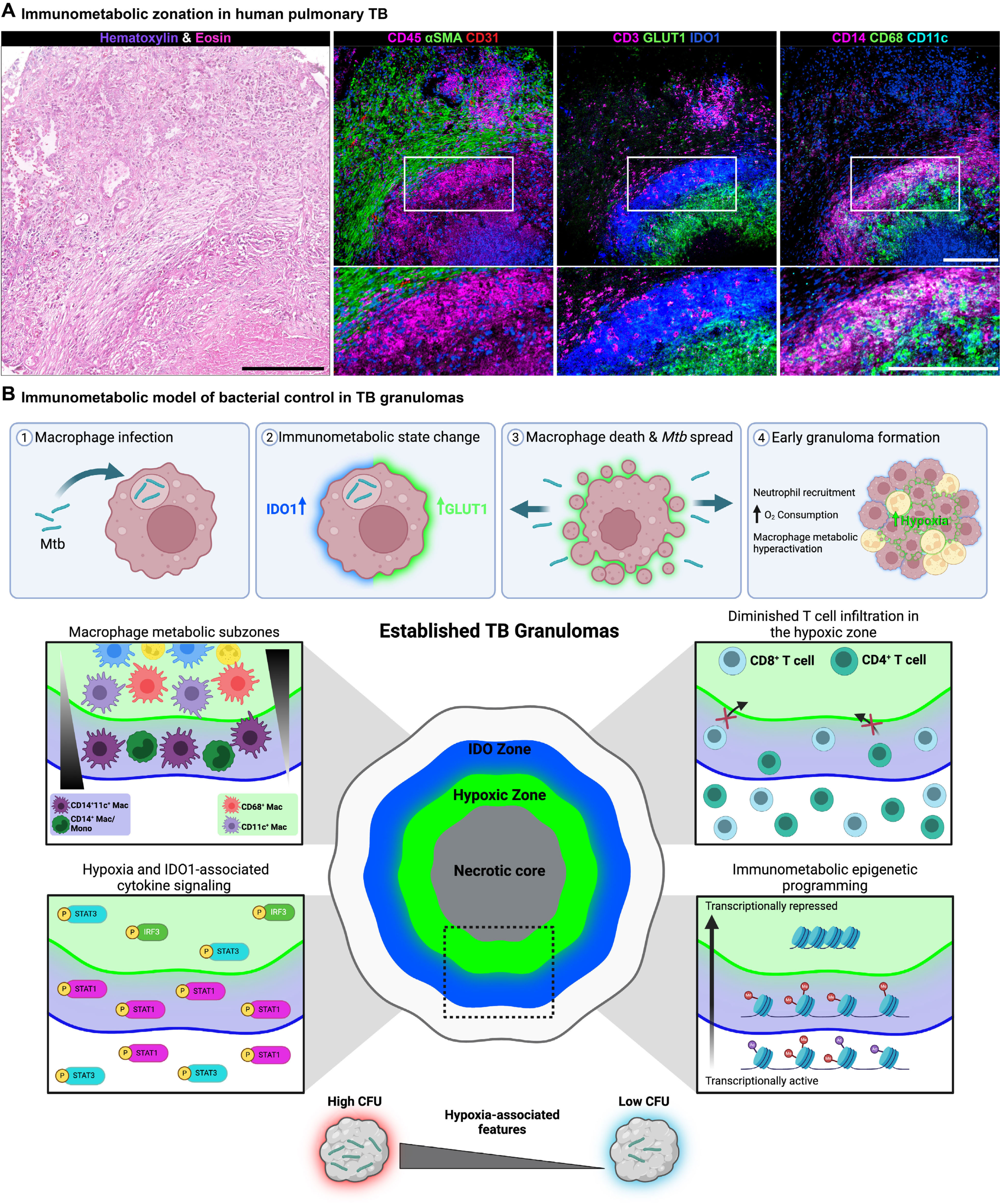
A model of immunometabolic regulation of granuloma organization and function in human TB. **A)** Representative images of a human pulmonary TB granuloma showing a serial section used for histological identification of the granuloma (left), CD45 (magenta), aSMA (green), CD31 (red), and HH3 (blue) (middle-left), CD3 (magenta), IDO1 (blue), and GLUT1 (green) (middle-right), and CD14 (magenta), CD68 (green), CD11c (cyan), and HH3 (blue) (right). Scale bar represents 200 μm. **B)** Schematic model of immunometabolic regulation of granuloma organization and function in human TB.

## DISCUSSION

The role of the TB granuloma has long been a subject of debate. While they can protect the host response by limiting intra-host bacterial spread and staging anti-Mtb immune responses, increasing evidence suggests they can also worsen the disease through excessive inflammation and function as a pro-bacterial haven. Ideal host immune responses fall into a ‘Goldilocks’ zone between pro- and anti-inflammatory signaling to achieve antibacterial responses that can clear Mtb without causing excessive tissue damage^6^. In our prior study examining archival human TB lesions, we observed the near ubiquitous immune-tolerant macrophage phenotype marked by expression of PD-L1, IDO1, TGFβ, and IL-10 and absence of IFNγ^8^. These macrophages were compartmentalized within the granuloma myeloid core together with Tregs. Notably, activated lymphocytes were preferentially depleted from this tolerogenic niche. Based on these findings and reports by others, we hypothesized that granuloma macrophages promote bacterial persistence by suppressing anti-Mtb immunity.

While the human TB granuloma study gave us a starting point for understanding myeloid-mediated immune tolerance during TB, the observational nature of examining archival human tissues imposes limitations on our ability to relate granuloma features with bacterial burden. Therefore, here we performed a comprehensive examination of TB lesions from Mtb-infected NHPs so that we could analyze tissue states where each granuloma has been characterized for its bacterial burden and time of development post infection. Access to this metadata allowed us to determine the microenvironmental features of the granuloma imprinting immunosuppressive qualities on macrophages, chart the relationship between granuloma macrophage and lymphocyte niches, and decode the contribution of these spatial properties to bacterial control during infection. In doing so, we identified hypoxia as a core driver of granuloma cellular organization and ability to support bacterial persistence. The central necrotic region of granulomas is associated with the presence of hypoxia, although little has been known about how this affects macrophage populations that are present in this region^17,50^. Here, we found that the myeloid core of granulomas bifurcates into two subzones. The first is the IDO zone, which is defined by high expression of IDO1, an enrichment of CD14-expressing macrophages and monocytes, and normoxia. In necrotic granulomas, a second subzone of the myeloid core appears. IDO1 expression drops off in this zone, while expression of GLUT1 sharply increases.

By referencing the spatial localization of GLUT1 with the *in-situ* hypoxia sensor, PIMO, we confirm that this secondary zone of the myeloid core is, indeed, hypoxic. This region was enriched for CD11c^+^ and CD68^+^ macrophages, both of which are associated with higher bacterial burden. The hypoxic zone also contained peri-necrotic neutrophils and a population of fibroblast-like FN1^+^ macrophages. After cross referencing these findings with scRNA-seq data, we found that hypoxic zone macrophages expressed *IL1RN*, *VEGFA*, and *IL6*, all of which have been associated with immunosuppression or otherwise dysfunctional immunity during TB^11,51,52^.

Using an algorithm inspired by GIS, we found evidence that the effects of hypoxia extend beyond the macrophage compartment and influence granuloma composition globally. While the outer IDO-expressing zone of the myeloid core exhibited a limited degree of infiltration, the hypoxic zone was virtually devoid of T-cells. While the absence of T cell migration into the myeloid core has been noted in TB granulomas and has implications for these cells’ ability to interact with Mtb-infected macrophages, reasons for their absence are not fully understood^53^. Our data suggest that hypoxia contributes to making this microenvironment inhospitable to T cell infiltration. Furthermore, macrophages in the hypoxic zone appeared to downregulate HLA-DR, which would limit antigen presentation to lymphocytes that do successfully infiltrate this environment. In this manner, hypoxia appears to create a topographical trap that inhibits T cell activity. In line with this we found that the cellular network in high bacterial burden granulomas predominately consists of hypoxia-associated spatial niches, suggesting that hypoxia is a global regulator of cellular organization and function in granulomas. Conversely, low bacterial burden granulomas were exemplified by increased cellular interactions with CD8^+^ T cells, endothelial cells, and IDO zone-enriched myeloid cells. Lastly, using archival tissue acquired from human patients with TB, we confirmed these architectural principles of NHP granuloma hypoxia are present in human lesions.

To understand how hypoxia establishes the tolerogenic environment in TB, we can look to the setting of solid tumors and leishmaniasis lesions, both of which exhibit strikingly similar hypoxia-imprinted microenvironments like we observe in TB granulomas. In solid tumors, especially in the setting of glioblastoma, hypoxia associates with suppressive tumor-associated macrophages and sequestration of tumor-infiltrating lymphocytes from tumor cells^25,26,30,54–56^. In these hypoxic tumor environments, lymphocytes cannot meet their metabolic demands, which include oxidative phosphorylation and glutamine metabolism^57,58^. This hypoxia-driven spatial patterning is termed ‘immune exclusion’ and closely matches the organization of hypoxic TB granulomas^19^. Similarly, during *Leishmania major* infection, hypoxia impairs the ability of macrophages to kill parasites via repression of iNOS activity, a microbiocidal pathway that has also been shown to mediate killing of Mtb by macrophages during murine and *in vitro* infections^59^. Both examples support our hypothesis that chronic hypoxia is a driver of suppressed immune cell activity and inefficient spatial patterning of adaptive immune responses.

Drawing on these examples and our findings, we have pieced together a theoretical timeline for the evolution of immunometabolic microenvironments in the TB granuloma. Our data suggests IDO1 is first expressed by granuloma macrophages. In non-caseating lesions, we observe that IDO1 and GLUT1 are co-expressed. This also points to the induction of aerobic glycolysis in activated macrophages, which is bolstered by the high uptake of the glucose analog ^18^F-FDG by both necrotic and non-necrotic TB granulomas during PET/CT imaging^60,61^. Furthermore, numerous studies demonstrate macrophages upregulate glycolysis upon infection to support bacterial killing and initiation of anti-bacterial immune responses^62–67^. While initially host-protective, this metabolic hyperactivation can lead to local oxygen depletion^68^. Furthermore, as a major virulence strategy, Mtb shunts cell death in macrophages from apoptosis to necrosis, which may seed the onset of hypoxia in a granuloma, prompting a feed-forward loop that drives increased levels of necrosis in oxygen-starved cells^69^. Neutrophils may also play a role in this pathology due to their swarming and subsequent NETosing in focal regions of the granuloma, and their high utilization of oxygen during inflammation^70,71^. This paired with sustained metabolic hyperactivation of infected macrophages and ongoing necrosis could synergize to rapidly deplete oxygen in a nascent TB granuloma. Ultimately in established granulomas, like those we analyzed in this study, this hypoxic milieu eventually partitions the myeloid core and excludes lymphocyte migration into regions associated with Mtb-infected macrophages

This model can provide insights into the immunoregulatory role of IDO1, an enzyme whose role in TB pathogenesis has been the subject of conflicting reports^42,72,73^. In our previous study we found that IDO1 expression was a universal attribute of TB granulomas that was not found in other granulomatous diseases, such as sarcoidosis^8,46^. It has been hypothesized that IDO1 inhibits immunity by promoting tolerance to Mtb and Treg activity. Consequently, inhibition of IDO1 is a candidate strategy for host-directed therapy for TB and has been shown to improve T cell infiltration of granulomas^42^. Here, we also find that IDO1 expression is ubiquitous, even in granulomas that sterilize infection, leading to questions about why this enzyme is so broadly expressed in granuloma macrophages. We found that IDO1 expression corresponded with evidence of type 1 interferon signaling, a transcriptionally repressed epigenetic state, and enrichment of high CFU-associated CD14^+^CD11c^+^ macrophages. In human TB granulomas, we previously observed these cells upregulate PD-L1^8^. Taken together, these characteristics suggest that IDO1 expression is a defining feature of a macrophage subset that can impair adaptive immunity. Alternatively, IDO1 may serve a host-protective role by restraining excessive inflammation and immune activation, thus further work into the mechanisms and consequences of IDO1 expression has important implications for understanding TB immunology.

While the IDO1’s role in granuloma function remains equivocal, our model strongly asserts that hypoxia is a keystone, pathological feature of the TB granuloma. We observe that hypoxia exerts its effects by promoting of a dysfunctional spatial design that suppresses innate and adaptive immune functions. Future explorations should refine the mechanisms by which hypoxia subverts host immunity to TB, including direct effects on expression of immune mediators that are regulated by HIF1 and indirect effects on the microenvironment, such as granuloma acidification or depletion of glutamine, which impair T cell effector functions and maturation^57,74^. However, hypoxia ultimately poses an obstacle to successful immune responses to Mtb and should be a high priority target for host-directed therapy. Strategies to ameliorate the effects of hypoxia include restoration of vascular integrity, direct delivery of oxygen to anoxic tissue environments, and breakdown of physical barriers to efficient oxygen diffusion such as fibrosis^75,76^. Several of these strategies have shown promise in reversing tumor hypoxia and even improving outcomes in preclinical animal models of TB^52,76–78^. Attempts to treat TB disease without addressing the physical and immunological barriers imposed by hypoxia will likely prove futile. Considering this, we propose that therapeutic approaches to promoting normoxia in TB granulomas may improve outcomes in TB by jumpstarting a cascade of host protective inflammation at the site of infection.

### Limitations of the Study

The significance of our findings should be contextualized within limitations of our study. Granulomas are first detectable by radiological imaging at 4-6 weeks post-infection with Mtb and the samples here represent 9-12 weeks post-infection. Therefore, we are limited in our interpretation of how the features we observed in our study manifest over time. Future studies would benefit from analysis of granulomas sampled across multiple timepoints of infection, with emphasis on early timepoints to validate our proposed model of the hypoxic milieu in TB granulomas. Our study cohort of granulomas were curated from archival tissue samples derived from multiple independent experiments. However, we did not observe any association between experimental batch and any of the central findings here. The central focus of our study was on how metabolic microenvironments in the granuloma associate with distinct cellular states and spatial patterning. Our MIBI-TOF antibody panel included only five metabolism-related proteins (CD36, Arginase-1, GLUT1, IDO1, and iNOS). By integrating scRNA-seq and spatial transcriptomics data into our study we were able to expand upon the immunometabolic relationships we identified with this limited set of proteins. Follow-on work using de novo mass spectrometry to directly quantify the spatial distribution of metabolites involved in these pathways could be informative and valuable. Lastly, due to the observational nature of our study, several important questions remain on the drivers of granuloma hypoxia and how perturbing hypoxia would alter granuloma structure and function. In future studies we will mechanistically assess these questions using in both *in vitro* organoid and *in vivo* animal models of TB granuloma formation that recapitulate the induction of hypoxia.

## METHODS

### Research Animals

This study leveraged archival formalin-fixed paraffin-embedded (FFPE) pulmonary tissue blocks from cynomolgus macaques *(Macaca fascicularis)* infected with low dose (2-13.5 CFU) *Mtb* Erdman via bronchoscopic instillation and followed until 9-12 weeks post-infection. Individual animal-level data can be found in **Table S1**.

### Study design and tissue cohort

#### Cynomolgus macaque TB granuloma cohort

FFPE *Mtb*-infected tissues were acquired from the University of Pittsburgh’s tissue repository from 16 cynomolgus macaques (n = 52 granulomas). Further information (including biological sex, number of granulomas, etc.) for each macaque involved in this study can be found in **Table S2**. All animal data for each specimen can be found in **Table S1.** Serial sections (5 μm) of each specimen were stained with hematoxylin-and-eosin (H&E) and inspected by an anatomic pathologist to screen for the presence of granulomatous inflammation. A wide variety of granuloma types were included: histologically solid granulomas, cellular granulomatous inflammation surrounding central necrosis, fibrotic granulomas, and multifocal granulomas. Regions of interest (ROIs) for imaging were chosen for each specimen based on the H&E images. No statistical methods were used to predetermine sample sizes.

### Cynomolgus macaque control tissues

In order to validate antibody reagents and optimize imaging settings we compiled a set of cynomolgus macaque control tissues from the University of Pittsburgh’s tissue repository. These included spleen, lymph node, lung, and colon specimens.

### Slide layout and construction

To minimize the impact of imaging batch effects, all cohort granuloma specimens were randomly arranged on 24 slides, with 1-3 specimens per slide. This prevented specimens from the same animal being placed on the same slide.

### Human TB granuloma

In collaboration with the Auria Biobank (Turku, Finland) we compiled a tissue microarray (TMA) of specimens from human patients with pulmonary TB. Working with a pathologist, we selected 0.5 mm diameter cores from each specimen to place on the TMA. A serial section (5 μm) of the TMA was stained with hematoxylin-and-eosin (H&E), while another section was analyzed via MIBI-TOF with an identical imaging panel as our NHP granuloma cohort.

### Immunofluorescence Staining

For detection of GLUT1, pimonidazole, and IDO1, we analyzed archival tissue from an animal that had been infected with Mtb for six weeks before being infused with pimonidazole for 48 hours and then necropsied. Formalin-fixed paraffin-embedded sections were mounted on superfrost-plus slides and antigen retrieval was performed as previously described^79^. Immunofluorescence staining was performed with anti-GLUT1 (rabbit polyclonal; #PA5-32428, Thermo Fisher Scientific, Waltham, MA), anti-pimonidazole (Mouse IgG1; Clone 4.3.11.3; Hydroxyprobe Inc, Burlington, MA) and anti-IDO1 (Mouse IgG1; clone OTI2G4; #LS-S-C174759, LifeSpan Biosciences). Species- and isotype-specific secondary antibodies for staining the anti-GLUT1 and anti-pimonidazole antibodies were purchased from Jackson ImmunoResearch Laboratories (West Grove, PA). IDO1 was detected with Thermo Fisher’s Zenon Mouse IgG1 Alexa Fluor 647 Labelling Kit (#Z25008, Thermo Fisher Scientific, Waltham, MA), which was used for tertiary staining. Another section in the same slide was stained in parallel and used as isotype control which received the same secondary staining to ensure specificity of each staining. Coverslips were mounted on the slides using ProLong Gold antifade reagent with DAPI (#P36931, Thermo Fisher Scientific). Images of granulomas were acquired at 20X using Nikon E1000 epifluorescence microscope with NIS elements AR 5.02.01. 64-bit software (Nikon Corporation, Melville, New York).

### MIBI-TOF Staining

#### Panel Construction

This is the first study to analyze cynomolgus macaque tissue with MIBI-TOF. Therefore, we had to develop macaque-compatible MIBI-TOF reagents. To achieve this, we first screened all reagents with chromogenic immunohistochemistry (IHC) to confirm epitope cross-reactivity and appropriate staining patterns in the control tissues described above (**Figure S2A**). All antibodies were then metal-labeled with the IonPath conjugation kit (IonPath, Menlo Park, USA) following the manufacturer instructions. To reduce reagent degradation and prolong shelf life, labeled antibodies were lyophilized in small batches and stored at -20°C. Antibodies were utilized for MIBI-TOF staining within 30 days of reconstitution. The appropriate antibody titer for MIBI-TOF was determined by staining control tissues with serial dilutions of each reagent between 0.0625-3 μg/mL. The final titer was chosen based on optimal signal-to-noise in these tissues. Information on all antibody reagents, metal reporters, and staining concentrations can be found in **Table S3**.

### Cohort Staining

Fresh aliquots of each antibody were reconstituted and tested within 72 hours of staining the tissue cohort. Antibodies were combined into a single master mix per the details found in **Table S3**. To avoid mistakes, each step in the staining protocol was performed in pairs, with one protocol reader and one pipettor. Due to performance issues with the Collagen-1 and pSTAT3 antibodies on a subset of slides, the signal from these channels was only analyzed on a subset of specimens. Complete details on all procedures related to reagent preparation, validation, and MIBI-TOF staining can found in the detailed protocols below:

### Interactive protocols

Reagent preparation: dx.doi.org/10.17504/protocols.io.bhmej43e

IHC staining: dx.doi.org/10.17504/protocols.io.bf6ajrae

MIBI staining: dx.doi.org/10.17504/protocols.io.dm6gprk2dvzp/v5

Sequenza staining: dx.doi.org/10.17504/protocols.io.bmc6k2ze

Antibody lyophilization: dx.doi.org/10.17504/protocols.io.bhmgj43w

Lyophilized antibody reconstitution: dx.doi.org/10.17504/protocols.io.bjd6ki9e

### MIBI-TOF data generation

All MIBI-TOF data was generated on a commercial MIBIScope instrument (IonPath, Menlo Park, USA). To image each ROI, we broke them down into 400 μm x 400 μm fields-of-view (FOVs). Each FOV was (1024 pixels)^2^ and were imaged using the “Coarse” settings on the MIBIScope instrument. Between all runs we conducted an imaging test on the molybdenum and PMMA standards per the manufacturer’s recommendations and settings. When the detector sensitivity dropped below the recommended levels, we adjusted the detector gain to minimize FOV-to-FOV sensitivity differences. One slide and one ROI was imaged at a time. However, to minimize image artifacts from instrument drift, we randomized the acquisition of each FOV within an ROI. All default image processing (background correction and denoising) were disabled.

### MIBI-TOF Data Low-level Processing

#### Data extraction

Low-level processing of the MIBI-TOF images was conducted with our publicly available, in-house pipeline, Toffy (https://github.com/angelolab/toffy). Overview of the low-level processing workflow and representative images of each step can be found in **Figure S3**. Following image extraction, we removed sources of signal contamination using a compensation method we developed called Rosetta. This approach implements channel-by-channel and pixel-by-pixel image compensation based on defined rates of signal contamination provided from the manufacturer (IonPath, Menlo Park, CA) and experimentally determined. The source and target channels, as well as the coefficients used for subtraction can be found in **Table S4**. Compensated images were then corrected for the instrument sensitivity drift that occurs throughout an imaging run. This was done using the median pulse height (MPH) normalization we developed and implemented in Toffy. Lastly, we observed that even with the steps applied above and the FOV randomization during imaging acquisition, the images displayed FOV-FOV non-uniformity in signal. To correct for this, we applied a tile-tile normalization approach. We used two channels for correction: HH3 for necrotic image regions and CD45 for non-necrotic image regions. For each FOV we determined the mean pixel value for the correction channel. Then we normalized each value to the max mean pixel value for the ROI. This normalized value was used a correction coefficient for each FOV. Per FOV, all image channels were multiplied by the correction coefficient with pixels in necrotic image regions and cellular necrotic regions differentially corrected.

### MIBI-TOF Data Analysis

#### Image region (necrosis, cellular, and bare slide) classification

Regions of necrosis were first identified by H&E-stained serial sections of each tissue. To annotate these regions, the HH3, CD45, and Vimentin images were overlaid and visualized in Fiji (ImageJ, version 2.9.0)^80^. The combination of these proteins allowed for visual identification of boundary between the cellular and necrotic regions of granulomas. The necrotic regions were manually annotated using the ROI Manager and freehand selection tools. The resulting annotations were exported as binary masks. Regions of bare slide background were automatically annotated by generating binary masks of the Au image channel. To produce this mask the Au signal per pixel was capped at 35 and smoothed with a Gaussian kernel (sigma = 2). Using the morphology toolkit from skimage (scikit-image, Python), asked regions with an area less than 2000 pixels were removed, and holes less than 1000 pixels were closed. Lastly, the mask was iteratively eroded five times. To cellular region of each specimen was defined as pixels not assigned to either the necrosis or bare slide masks. Example masks can be found in **Figure S4**.

### Adjusting for corrupted fields of view

During image acquisition, 56 FOVs (<4% of total, randomly distributed across FOVs) were corrupted during the transfer of data from the MIBIScope to the raw data file due to a software bug. Due to file corruption, the FOVs are left blank where they appear in the ROIs. All masks were corrected for this to ensure area-based calculations or normalizations were accurate.

### Single cell segmentation

Using a publicly available image analysis pipeline developed by our group, ark-analysis (https://github.com/angelolab/ark-analysis), we used Mesmer to segment individual cells from all images. We combined CD45 and CD14 to form a single membrane channel and HH3, H3K9Ac, and H3K27me3 to form a single nuclear channel. We used the default Mesmer parameters with one exception. To improve the accuracy of macrophage segmentation, we modified to interior threshold for the watershed step to 0.2. Cell segmentation was conducted on individual FOVs (as opposed to ROIs) separately and then combined into a single segmentation mask for each ROI.

### Giant cell segmentation

Multinucleated giant cells were manually segmented. They were first identified in H&E-stained serial sections of each tissue. To identify them in the MIBI-TOF images CD45, HH3, CD68, and CHIT1 were overlaid and visualized in Fiji (ImageJ, version 2.9.0). The ROI Manager and freehand selection tools were used to annotate their borders. Binary masks for each giant cell were exported and then appended to the Mesmer-derived segmentation masks. Any Mesmer-derived cell objects that overlapped with the giant cell regions were removed from the final segmentation masks.

### Pixel and cell clustering

Using ark-analysis, we conducted pixel clustering was performed using Pixie, a clustering algorithm optimized for multiplexed image data. We used the following parameters: blur_factor = 2 and subset_proportion = 0.1. We used the CD3, CD4, CD8a, CD11c, CD14, CD20, CD31, CD45, CD68, Vimentin, CD163, CD206, Calprotectin, Chymase/Tryptase, αSMA, DC-LAMP, Fibronectin, HLA-DR, and CD36 channels for clustering. We initially over-clustered the data into 100 clusters which were then combined into 22 meta-clusters (**Figure S5**). We next determined the proportion of each pixel cluster in each segmented cell object. These profiles were then used to cluster cells, first over clustered into 100 clusters and then combined into 20 metaclusters. To confirm accurate clustering and correct system instances of inaccurate assignment we inspected the clusters in Mantis Viewer. We then corrected cell objects that had been misassigned by reassigning their phenotypic assignment. Giant cells were manually assigned their phenotype based on the previously described giant cell segmentation approach.

### Metabolic zone annotation

To generate the IDO zone masks, the IDO1 signal from each specimen was filtered with a threshold to remove noise, smoothed with a Gaussian blur, and further filtered to remove holes and small objects. The resulting masks were exported as binary .tiff files. To generate the hypoxic zone masks, we utilized the GLUT1 channel. However, to exclude the regions were IDO1 and GLUT1 co-localize in the IDO zone, we removed the GLUT1 signal that corresponded with IDO1 signal. The filtered GLUT1 signal from each specimen was further processed with a threshold to remove noise, smoothed with a Gaussian blur, and further filtered to remove holes and small objects. The resulting masks were exported as binary .tiff files.

### CFU-based analyses

For each specimen we determined the frequency of each cell phenotype out of total cells. We performed cellular correlation analyses using a Spearman correlation calculation (rcorr function, Hmsic, R package) and then corrected the resulting p-values with a false discovery rate of 5% (p.adjust function, R). We also bifurcated samples into ‘high’ and ‘low’ CFU groups based on the median log_10_ CFU per specimen (3.26).

### Metabolic zone enrichment

We first combined the cell phenotype maps and metabolic zone masks to assign cells to either the IDO or hypoxic zone. To calculate the zone enrichment score we enumerated the count of each cell phenotype in either the IDO or hypoxic zone. If a cell object’s area overlapped with both zones, the tie break was based on which zone had a greater proportion of that cell’s area. In the event of a true tie (19 / 392659 cells) the cell was assigned to the hypoxic zone. Next, per specimen, we normalized the count of each cell phenotype by the zone area. To compute the enrichment score we took the log_2_ fold-change of the cell phenotype density in the hypoxic zone to the cell phenotype density in the IDO zone. For the myeloid core enrichment shown in **Figure S7A** we performed the same calculation except the cells were classified based on whether they localized to the myeloid core (IDO and hypoxic zones combined).

### T cell-macrophage association score

To evaluate the spatial association between macrophages and T cells we used our previously developed ‘mixing score’ method. Briefly, for each ROI and for each zone we determined the number of times T cells (CD4^+^ or CD8^+^ T cell) were within 10 μm of macrophages (CD11c^+^ Macs, CD14^+^CD11c^+^ Macs, CD68^+^ Macs, CD14^+^ Mac/Monos, CD163^+^ Macs, CD206^+^ Macs, FN1^+^ Macs, and multinucleated giant cells). This value was divided by the number of times macrophages were within 10 μm of other macrophages. In addition to performing this analysis in bulk, we also broke it down to evaluate CD4^+^ or CD8^+^ T cell associations separately.

### Immunotopography (buffer) analysis

In order to perform buffer analysis, we first created binary masks defining the centroids of each granuloma. For necrotic regions we used the previously generated necrosis masks. For regions with solid granulomas, we manually defined the granuloma centroid based on the IDO1 and CD14 images and placed a centroid dot with a radius = 20 pixels. Each mask was also padded with 2048 pixels in the x and y directions for buffer expansion. Binary masks were imported to ArcGIS Pro (Version 3.2, Esri, Redlands, CA) as rasters. The input raster images were reclassified into polygons. Next the “Multiple Ring Buffer” tool was used with the “geodesic” method option to generate concentric buffer rings around each granuloma center polygon. A ring size of ‘10 meters’ was used which corresponded to a ring thickness of ∼60-90 pixels depending on the image. The number of buffer rings was determined empirically based on the maximum number of buffer rings that could be appended until the image boundaries (indicated by the image padding) were reached. Resulting buffer masks were exported as .tif files. Using the skimage and scipy libraries in Python the padding was removed, and the buffer mask was converted to a labeled mask where buffers were labeled 1→n with 1 being assigned to the innermost buffer and n representing the outermost buffer. To extract cellular, protein, and metabolic zone feature from each buffer, the buffer mask was combined with the cellular mask exclude background signal from areas of bare slide. For each specimen and for each buffer, the mean pixel intensity of each protein was quantified, as well as the density (area-normalized count) of each cell type from the cell phenotype map. A cell was assigned to a buffer zone if at least 50% of its area overlapped with that zone. The metabolic zone maps for the IDO and hypoxic zones were combined with the buffer mask to enumerate the proportion of each buffer zone that was assigned to each metabolic zone. To integrate the buffer-derived data across all specimens the buffer zone numbers were normalized to the total number of buffers per specimen. These values were then partitioned into 20 bins.

### QUantitative InterCellular nicHe Enrichment (QUICHE)

To test for differences in cellular organization across conditions (i.e., high vs low CFU), we performed spatial statistical differential abundance testing using QUICHE (https://github.com/jranek/quiche) in a three-step process^49^. Briefly, we identified local cellular niches within each image according to spatial proximity (*radius = 200 pixels*), performed distribution-focused downsampling to select a subset of niches (*niches = 2500*) from each sample for condition comparisons, and constructed a niche similarity graph for differential enrichment testing (*k_sim = 100*). QUICHE niche neighborhoods were considered significant if spatial false discovery rate (FDR) < 0.05^81^.

### Radial Comparison for Neutrophils

Using the necrosis masks, for each necrosis object, neutrophils were annotated if their centroid fell within a 25 μm boundary from the necrosis border; otherwise, neutrophils were annotated as outside the boundary. Neutrophils were grouped by location with respect to the boundary, and the log2 fold change was calculated between outer boundary cells and inner boundary cells for each sample. A Wilcoxon signed-rank test was utilized to statistically compare the distribution of log2 fold changes between high and low CFU samples.

### Differential Expression within Necrosis

Using the necrosis masks, marker expression was summed across the entire necrosis area for each necrosis region per sample. The sum of the expression per necrosis region was then normalized by its area. Then, the normalized marker expression per necrosis object was summed to generate a normalized necrosis marker expression value per sample. The log2 fold change was calculated between the high and low burden samples. A Student’s t-test was applied to statistically compare the difference in normalized expression between the high and low burden samples. The results for each marker were plotted as a volcano plot.

### Single-cell RNA sequencing analysis

The scRNA-seq dataset of NHP granulomas analyzed here in published in Peters et al^46^. In order to identify clusters most closely matching those identified with multiplexed imaging we evaluated expression of *IDO1*, *SLC2A1*, and *SLC2A3*. From this we determined the originally annotated clusters Recruited Macrophage 2 (RM2) and Recruited Macrophage 3 (RM3) as Hypoxic Zone Macs and IDO Zone Macs, respectively. The Glycolysis/Gluconeogenesis Score was evaluated using scMetabolism (scMetabolism, R package). Differential gene expression was determined based on log-fold change of expression between clusters with p-values determined with a Wilcoxon rank sum test and adjusted with Bonferroni correction (FindMarkers function in Seurat, R package). Gene set enrichment analysis (GSEA) was performed to evaluate differential pathway utilization between clusters using Enrichr (enrichR, R package).

### Spatial transcriptomics

The Visium CytAssist Spatial Gene Expression Reagent kit, 6.5 mm, human, was used to process a serial section of NHP6-Granuloma 3 following the 10X Genomics User Guide CG000495, Rev E (10X Genomics, Pleasanton, CA). Briefly, tissue sections were H&E stained and imaged on the Zeiss AxioScan microscope, and then processed according to the user guide. The cycle number determination was performed for each sample and final libraries were amplified using the Dual Index TS Set A kit, following standard Illumina multiplexing guidelines. Initially, to assess if the human probes sufficiently detected NHP tissue, library samples were prepared, quantified using the Kapa Library Quantification kit (Roche Sequencing Solutions, Indianapolis, IN), and sequenced at low depth on a MiSeq instrument run as 28 X 50 bp paired- end reads using the Micro kit, v2, 300 cycle chemistries (Illumina, San Diego, CA). For the deep sequencing, sample libraries were quantified using the 2100 Bioanalyzer, pooled using adjusted volumes according to approximate spot coverage, and sequenced on the NextSeq 2000 instrument as 28 X 50 bp paired-end reads using the P3, 100 cycle kit (Illumina, San Diego, CA). Raw sequence files were processed through the SpaceRanger v2.0.0. (BCLs) were demultiplexed into fastq files using SpaceRanger mkfastq. The resulting fastq.gz files were used as input into SpaceRanger count using the Visium Human Transcriptome probe set v2.0 GRCh38 as a reference. Image data from the H&E stained and CytAssist instrument files were also used in the SpaceRanger CytAssist-enabled workflow. Visualization of transcript expression across spots was done using the SpatialFeaturePlot function in Seurat version 5.1.0.

### Figure generation

Figures were produced using Adobe Illustrator, Adobe Photoshop, and BioRender.

### Data availability

All images, segmentation masks, and single cell data will be made available upon publication in a peer-reviewed journal. For access prior to publication please contact the corresponding author.

### Code availability

All original code used to perform analyses and generate figures will be made publicly available upon publication in a peer-reviewed journal. For access prior to publication please contact the corresponding author.

## ACKNOWLEDGMENTS

We would like to thank Pauline Chu, the Stanford Human Histology Core, and Jim Cherry, Daniel Bruno, and Elizabeth Fischer of the NIAID Research Technologies Branch for their assistance with this study. We would also like to thank Adrie Steyn, Alex Shalek, and Daniel Barber for insightful discussions related to this work. We acknowledge the Auria Biobank for providing the human TB tissue specimens. This study was supported by the Bill & Melinda Gates Foundation and the Wellcome Leap Delta Tissue program. E.F.M., S.D., S.A., C.M., and S.W. received support from the National Institutes of Health Division of Intramural Research. N.F.G. was supported by NCI CA246880, NCI CA264307, and the Stanford Graduate Fellowship. B.B. was supported by the National Institutes of Health (R01AI166313 and R01AI164970). J.M was supported by the National Institutes of Health (AI164970 and AI164970).

## AUTHOR CONTRIBUTIONS

E.F.M., M.A., J.L.F. J.T.M., S.F., and D.K. conceived and designed the study; J.L.F., J.T.M., P.L.L., C.S., M.A.R., E.K., P.M., and C.L.B. curated the NHP tissue specimens and all associated metadata; R.V. and J.T.M. analyzed the pimonidazole-labeled granuloma; E.F.M., A.D., C.C.F., M.G., P.B., and M.B. performed reagent optimization and generated the MIBI-TOF data. E.F.M., A.D., J.S.R., C.C.L., N.F.G., Y.B., C.S., and A.K. processed and analyzed the MIBI data; J.P. and B.D.B. conducted the scRNA sequencing data analysis. E.F.M., I.F., C.G., and J.R.B. developed the immunotopography analysis pipeline; S.A., C.M., S.W., and S.D. conducted the spatial transcriptomics analysis. E.F.M. and A.D. wrote the manuscript. E.F.M. and A.D. generated the figures. J.T.M., J.L.F., and M.A. supervised the project. All authors provided feedback on the manuscript.

---

**Figure S1:**
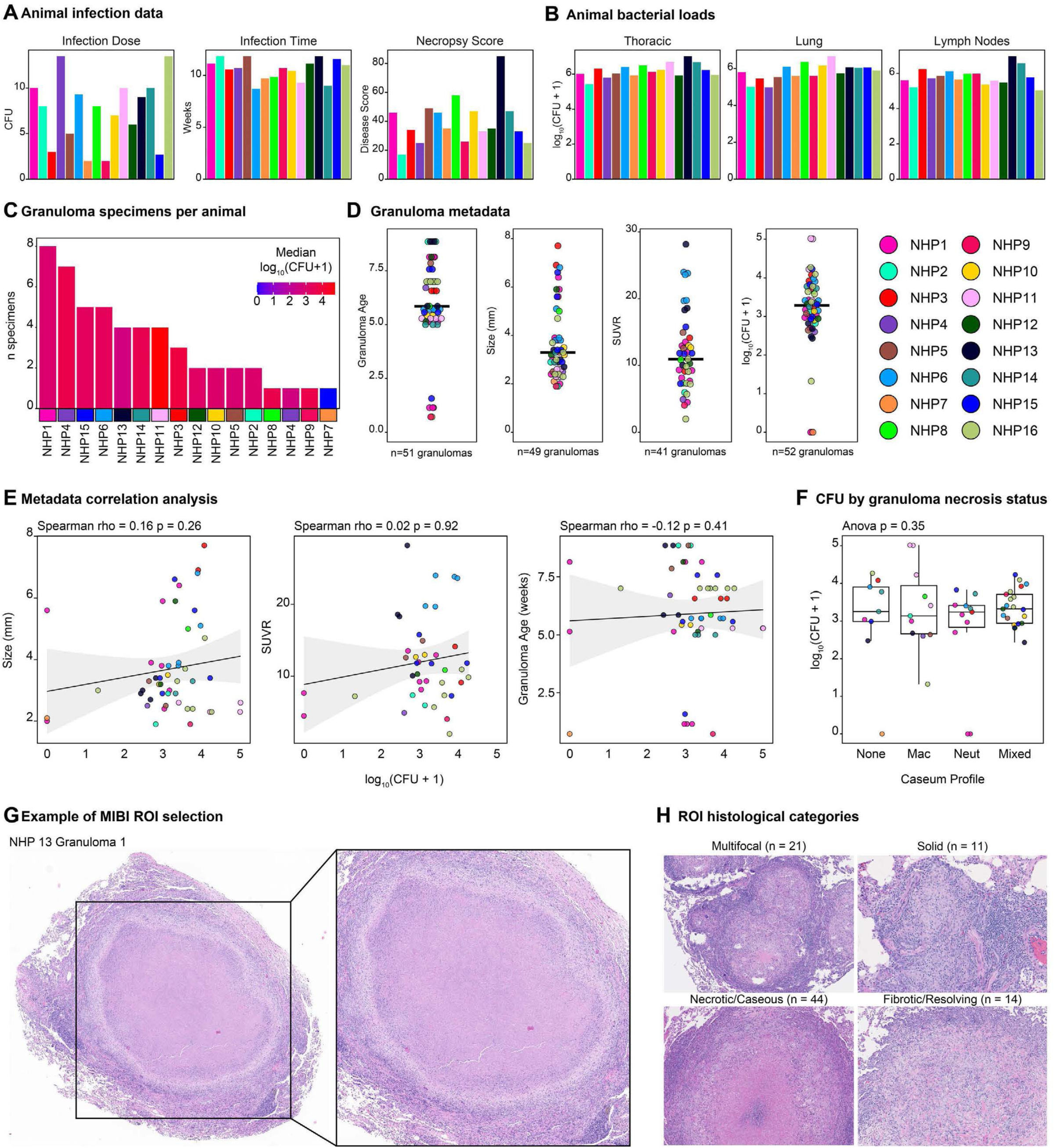
NHP TB granuloma cohort attributes. **A)** Infection data for study animals including infection inoculum dose (left), infection time (middle), and necropsy score (right). **B)** Bulk bacterial loads (log_10_[CFU+1]) for the thoracic area (left), lungs (middle), and lymph nodes (right). **C)** Number of granuloma specimens per animal in descending order. Fill indicates the median CFU for the specimens from that animal. **D)** The age (left), size (left-middle), PET probe uptake (SUVR, right-middle), and bacterial burden (log_10_[CFU+1], right) per granuloma. Each dot represents one granuloma, and the color represents the animal from which the granuloma was derived. **E)** Bi-axial scatter plot of granuloma bacterial burden versus size (left), PET probe uptake (middle), and age (right). Linear regression (black solid line) with 95% confidence interval (CI; grey silhouette). Spearman rho and p-value (two-tailed t test) for each relationship is displayed above the plot. **F)** Bacterial burden of granulomas broken down by caseum type. *P* value was determined using ANOVA. Boxplots display the median and IQR (25–75%), with whiskers representing the upper- and lower-quartile ±1.5× IQR. **G)** Example of a MIBI-TOF imaging region selected from a hematoxylin and eosin-stained NHP TB granuloma. H) Breakdown of TB granuloma cohort by histological category.

**Figure S2:**
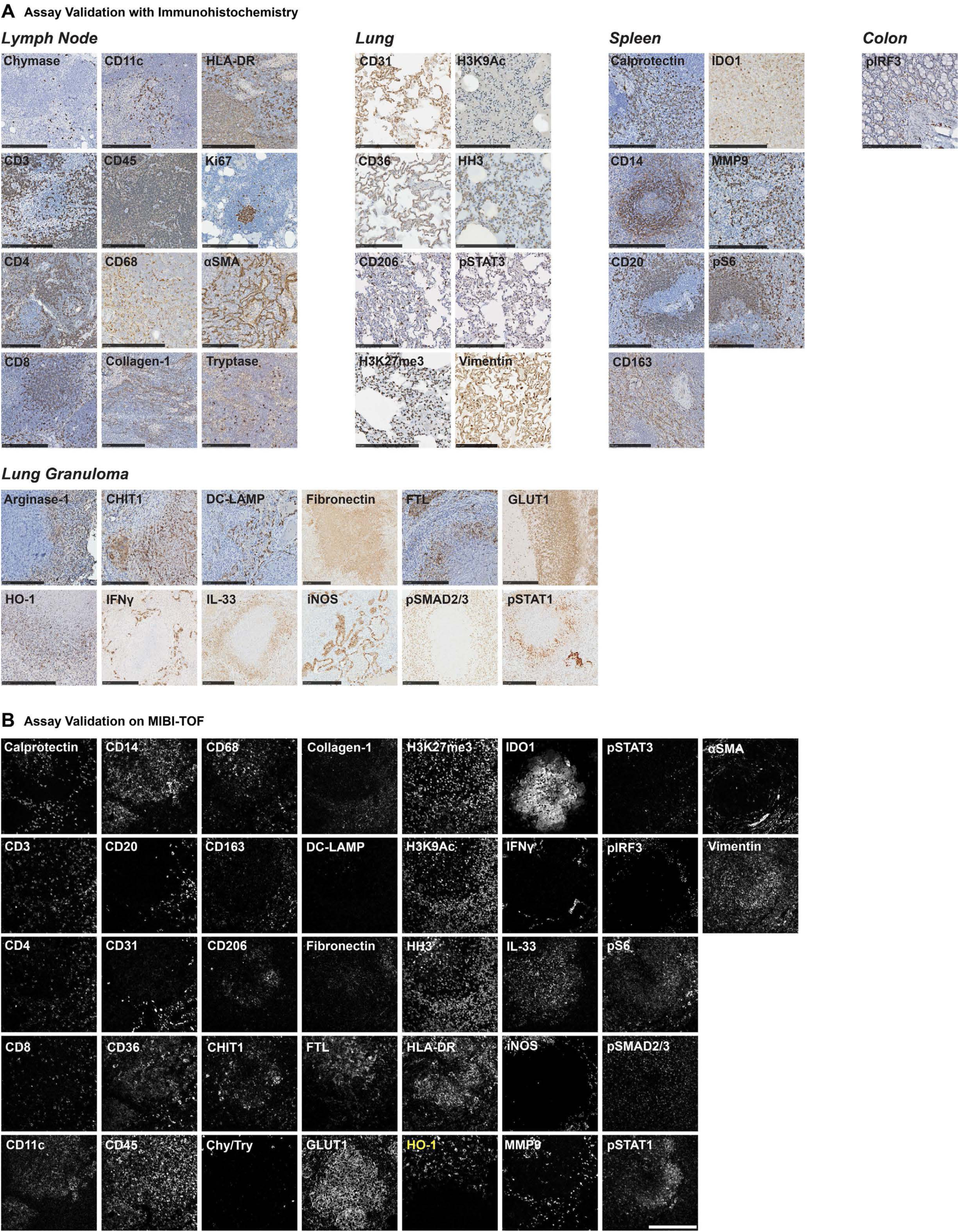
Validation of NHP-reactive antibody reagents. **A)** Representative chromogenic immunohistochemistry images for all antibody targets in lymph node, lung, spleen, colon, or granuloma tissue. **B)** Representative MIBI-TOF images for all antibody targets in NHP TB granulomas. Representative image of HO-1 (yellow text) comes from rhesus macaque granulomatous tissue. Scale bar represents 200 μm.

**Figure S3:**
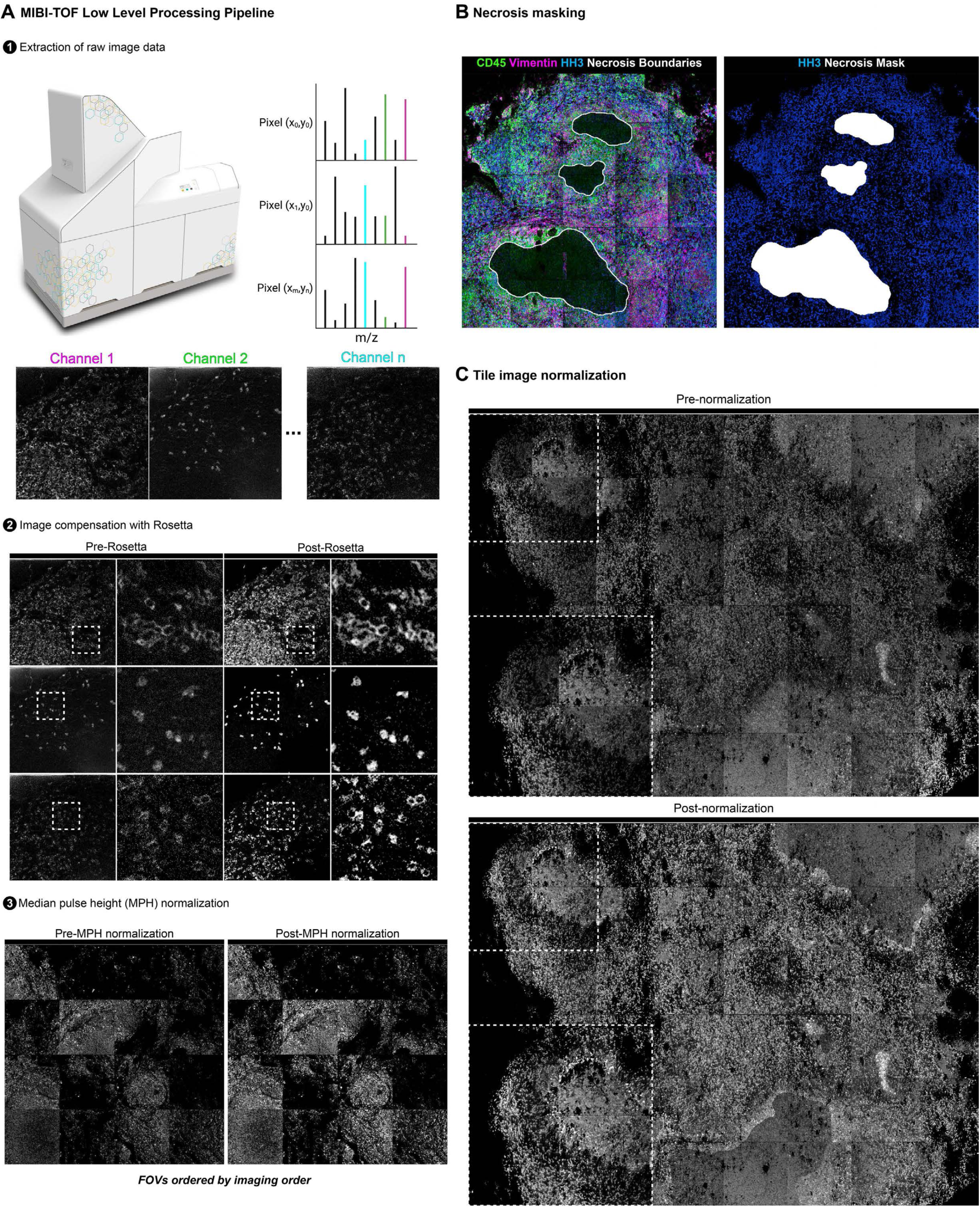
Low-level MIBI-TOF data processing pipeline. **A)** Overview of MIBI-TOF low level processing pipeline for data extraction (top), image compensation (middle), and normalization (bottom). Each stage displays representative images before and after each stage of processing. **B)** Example of necrosis masking procedure with the composite image used to identify necrotic regions (left, green = CD45, magenta = Vimentin, blue = nuclei, white = necrosis boundaries) and the resulting masked regions (right, blue = nuclei, white = necrotic zones). **C)** Representative image channel before (top) and after (bottom) tile-tile normalization.

**Figure S4:**
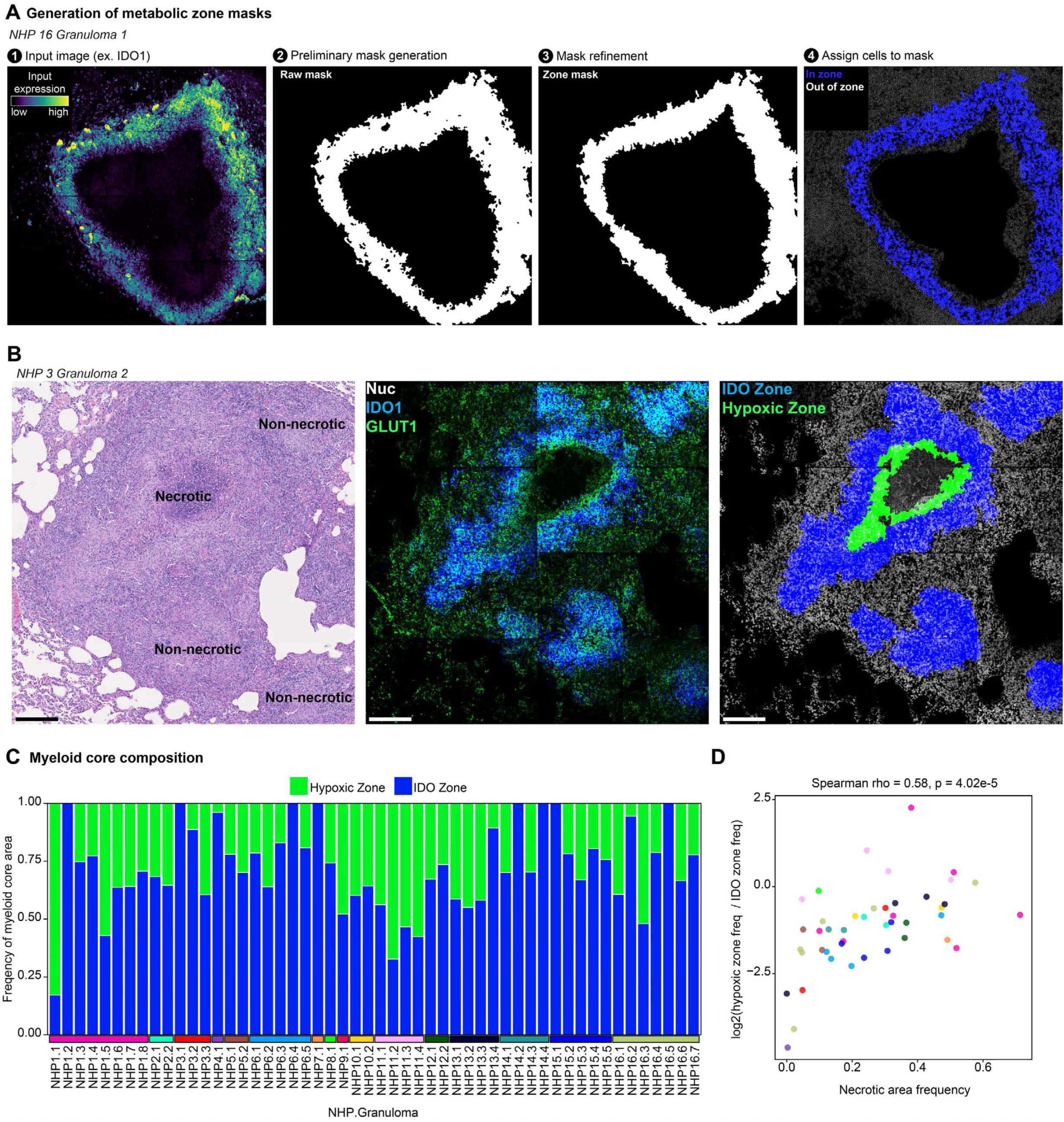
Delineating granuloma metabolic zonation. **A)** Overview of the metabolic zone masking process for representative granuloma showing (from left to right) the raw signal, unprocessed mask, closed mask, and annotation of segmented cells based on metabolic zone assignment. **B)** Representative granuloma with necrotic and non-necrotic regions annotated via histology (left), raw metabolic signal (center, green = GLUT1, blue = IDO1), and the annotated metabolic masks (right, blue = IDO zone, green = GLUT1 zone, white = nuclei). Scale bar represents 200 μm. **C)** Myeloid core area broken down by composition. **D)** Ratio of myeloid core area composition against necrotic area frequency. Dots represent individual granulomas and are colored by animal. Spearman rho and p-value (two-tailed t test) is displayed above the plot.

**Figure S5:**
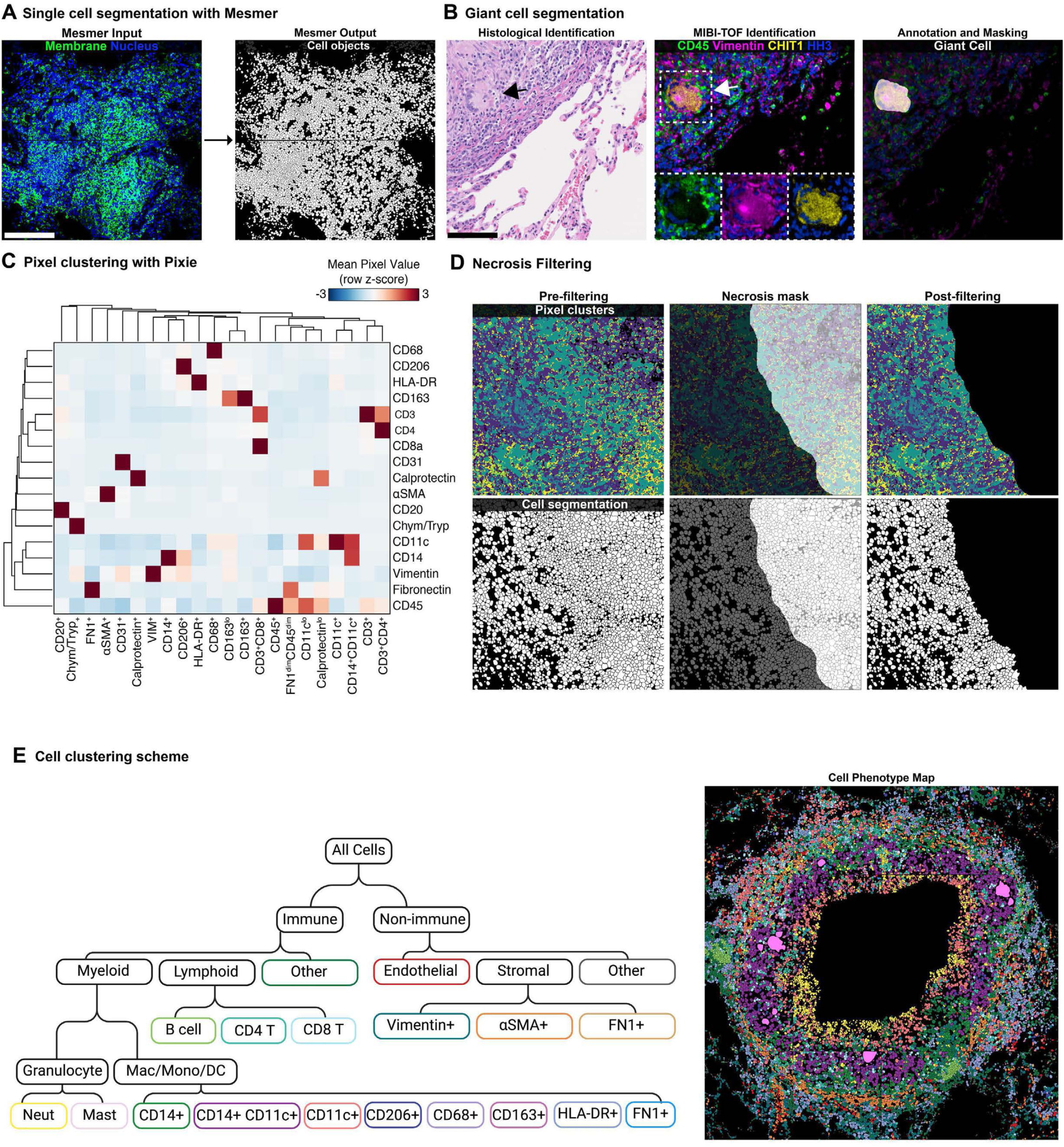
Enumerating and phenotyping single cells from MIBI-TOF data. **A)** Overview of Mesmer single cell segmentation pipeline showing a representative input image (green = membrane signal, blue = nuclear signal) and output (cell object = white). Scale bar represents 200 μm. **B)** Example of multinucleated giant cell segmentation showing identification of a giant cell via histology (left) and MIBI-TOF (middle, green = CD45, magenta = Vimentin, yellow = CHIT1, blue = nuclei), and the resulting giant cell segmentation mask (right). **C)** Pixel clusters defined with Pixie. Heatmap shows the mean value of each image channel (rows, row z-scored) in each pixel cluster (columns). Columns and rows are hierarchically clustered (complete linkage). **D)** Example of necrosis-filtering of pixel clusters and segmented cell objects. **E)** Conceptual overview of cell phenotyping strategy (left) and a representative cell phenotype where segmented cell objects are colored by their phenotype.

**Figure S6:**
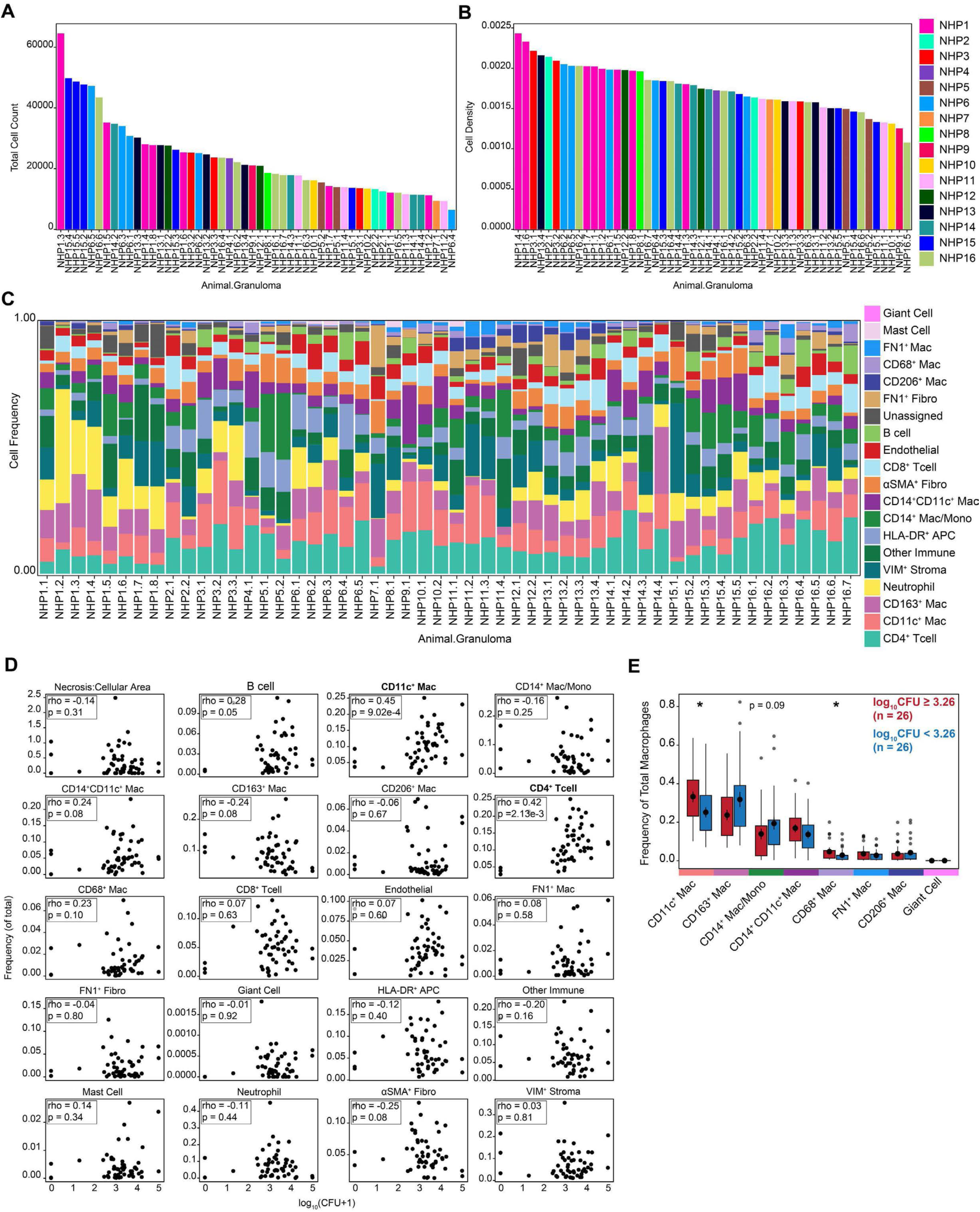
Analysis of granuloma cellular composition. **A)** The total number of cells per granuloma specimen in descending order. Color indicates the animal. **B)** The area-normalized cellular density per granuloma specimen in descending order. Color indicates the animal. **C)** The single cell composition of each granuloma. Cell phenotypes were ordered by descending median frequency. **D)** The frequency of each cell type (of all cells) versus the bacterial burden of each granuloma. Each plot displays the Spearman rho and p-value (two-tailed t test). The header is bolded for relationships with statistical significance. **E)** The frequency of each macrophage subset (of total macrophages) between granulomas with high (red) versus low (blue) bacterial burden. Macrophage populations are ordered by descending median frequency overall. Dots represent the mean and standard error. Boxplots display the median and IQR (25–75%), with whiskers representing the upper- and lower-quartile ±1.5× IQR. *P* values were calculated with a Wilcoxon rank-sum test (two tailed) (*P < 0.05).

**Figure S7:**
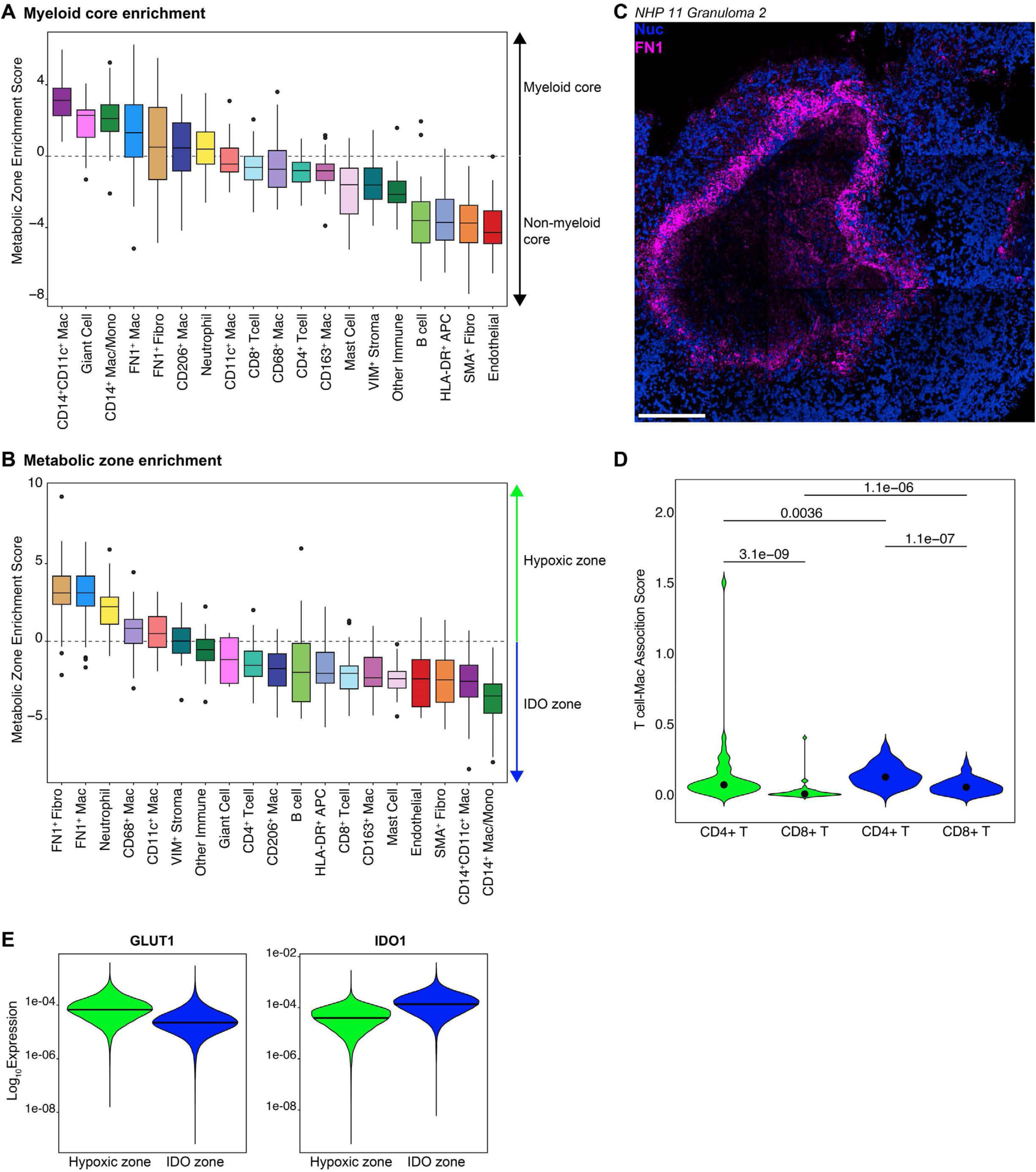
Immunometabolic distribution of cells. **A)** Myeloid core enrichment of granuloma immune and non-immune cells. Boxes are ordered by descending median and colored by cell type. **B)** Metabolic zone enrichment of myeloid-core infiltrating cells. Boxes are ordered by descending median and colored by cell type. Boxplots display the median and IQR (25–75%), with whiskers representing the upper- and lower-quartile ±1.5× IQR. **C)** Representative image of FN1-expressing (magenta, blue = nuclei) macrophages in the granuloma myeloid core. Scale bar represents 200 μm. **D)** T cell-macrophage association score in the hypoxic zone (green) and IDO zone (blue). Dot represents the mean. *P* values were calculated with a Wilcoxon rank-sum test (two tailed). **E)** Log_10_-normalized expression of GLUT1 (left) and IDO1 (right) in all macrophages from the hypoxic (green) or IDO (blue) zones.

**Figure S8:**
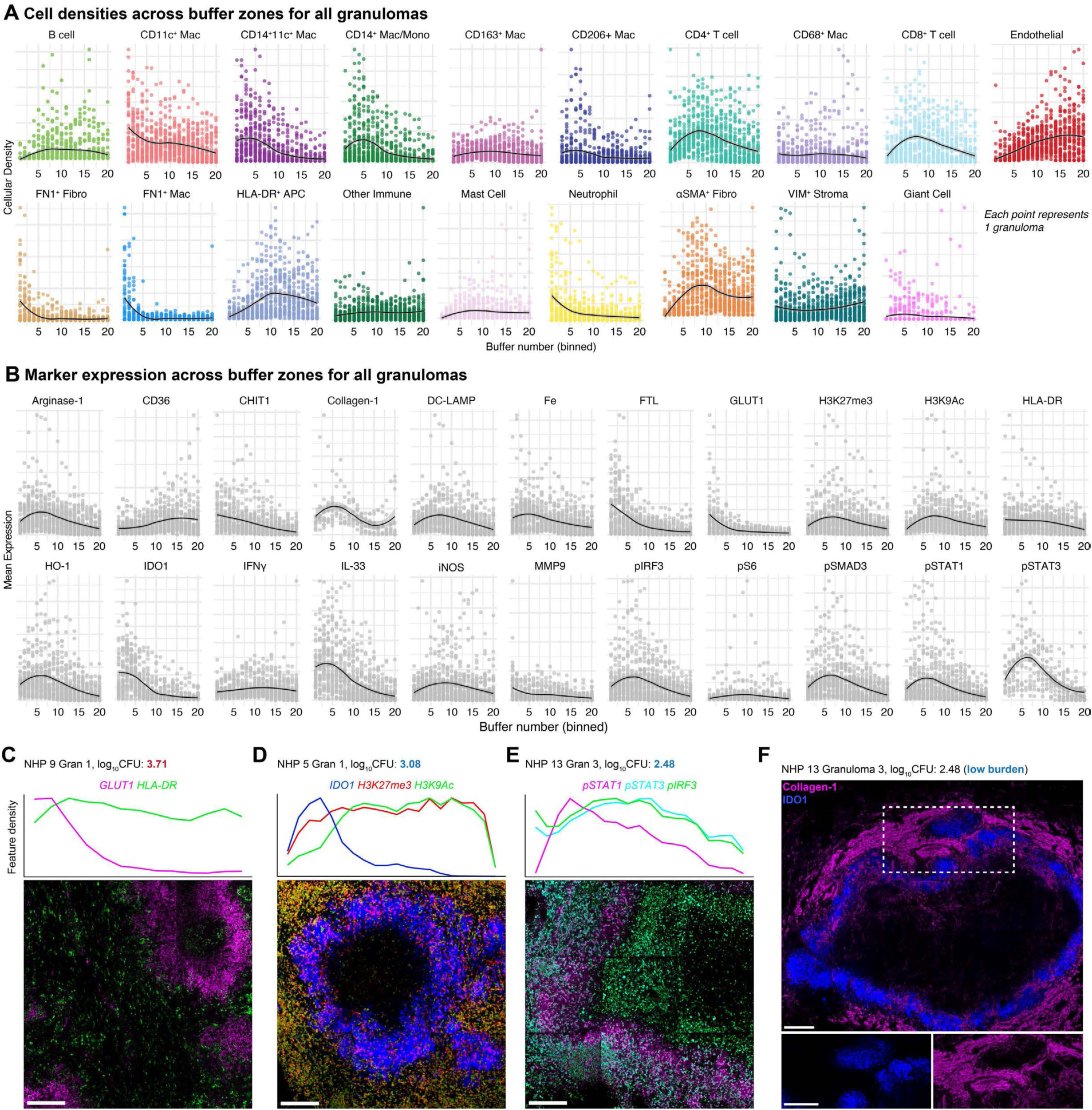
Radial immunotopography of granulomas. **A)** Cellular densities across binned buffer zones for all granulomas. Each point represents one granuloma. Black fitted line displays the local polynomial regression with the gray silhouette indicating the 95% confidence interval. **B)** Mean expression of each of protein across binned buffer zones for all granulomas. Each point represents one granuloma. Black fitted line displays the local polynomial regression with the gray silhouette indicating the 95% confidence interval. **C-E)** Track plot (top) and representative image of the radial distribution of **C)** GLUT1 (magenta) and HLA-DR (green) in NHP 9 granuloma 1, **D)** IDO1 (blue), H3K27me3 (red), and H3K9Ac (green) in NHP 6 granuloma 1, and **E)** pSTAT1 (magenta), pSTAT3 (cyan), and pIRF3 (green) in NHP 13 granuloma 3. **F)** Representative image of collagen-1 (magenta) and IDO1 (blue) with zoomed inset (white dashed box) displayed on the right. Scale bars represent 200 μm.

**Figure S9:**
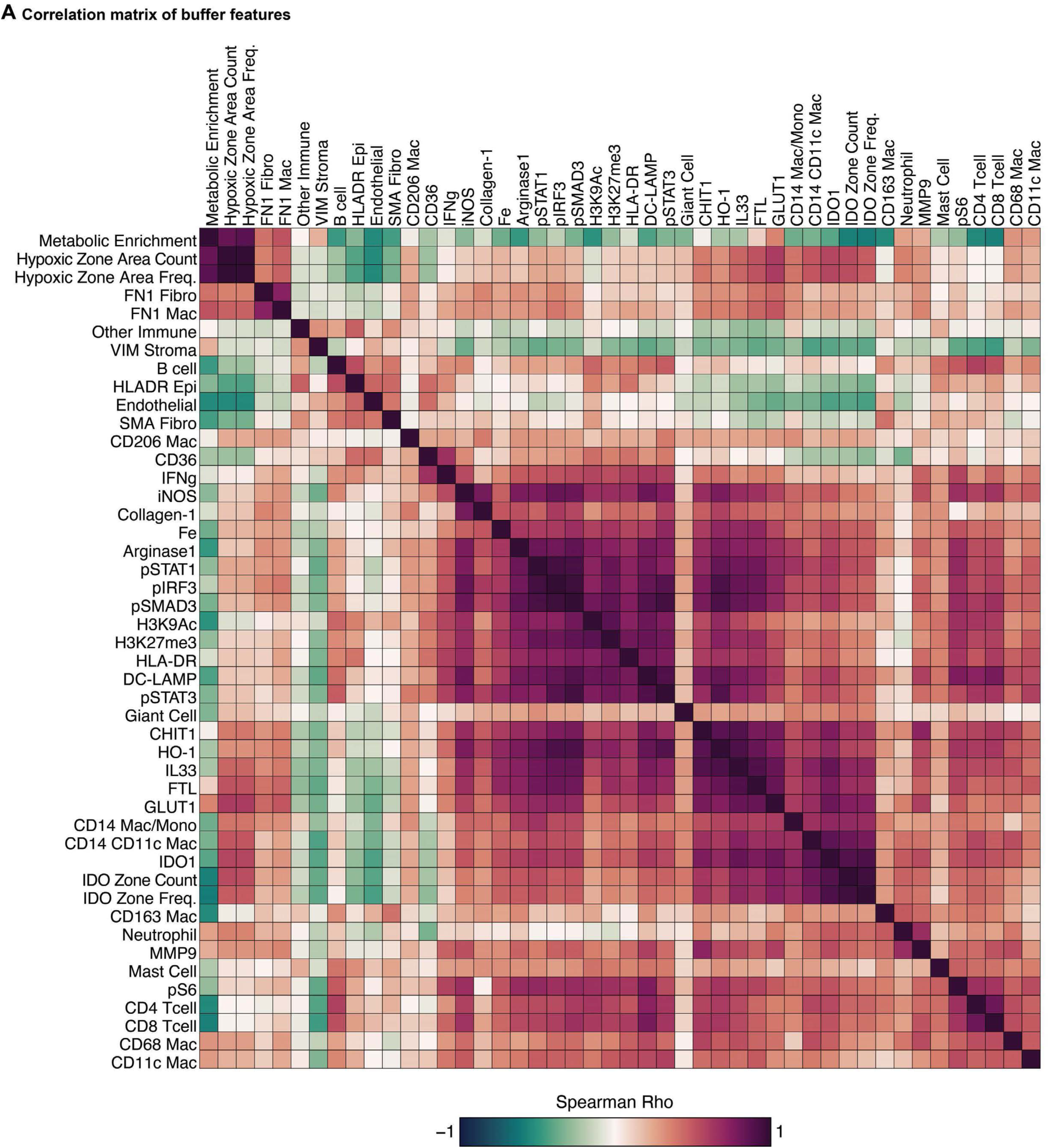
Buffer-based correlation analysis. Spearman correlation for all cellular, protein, and metabolic buffers on a buffer-by-buffer basis.

**Figure S10:**
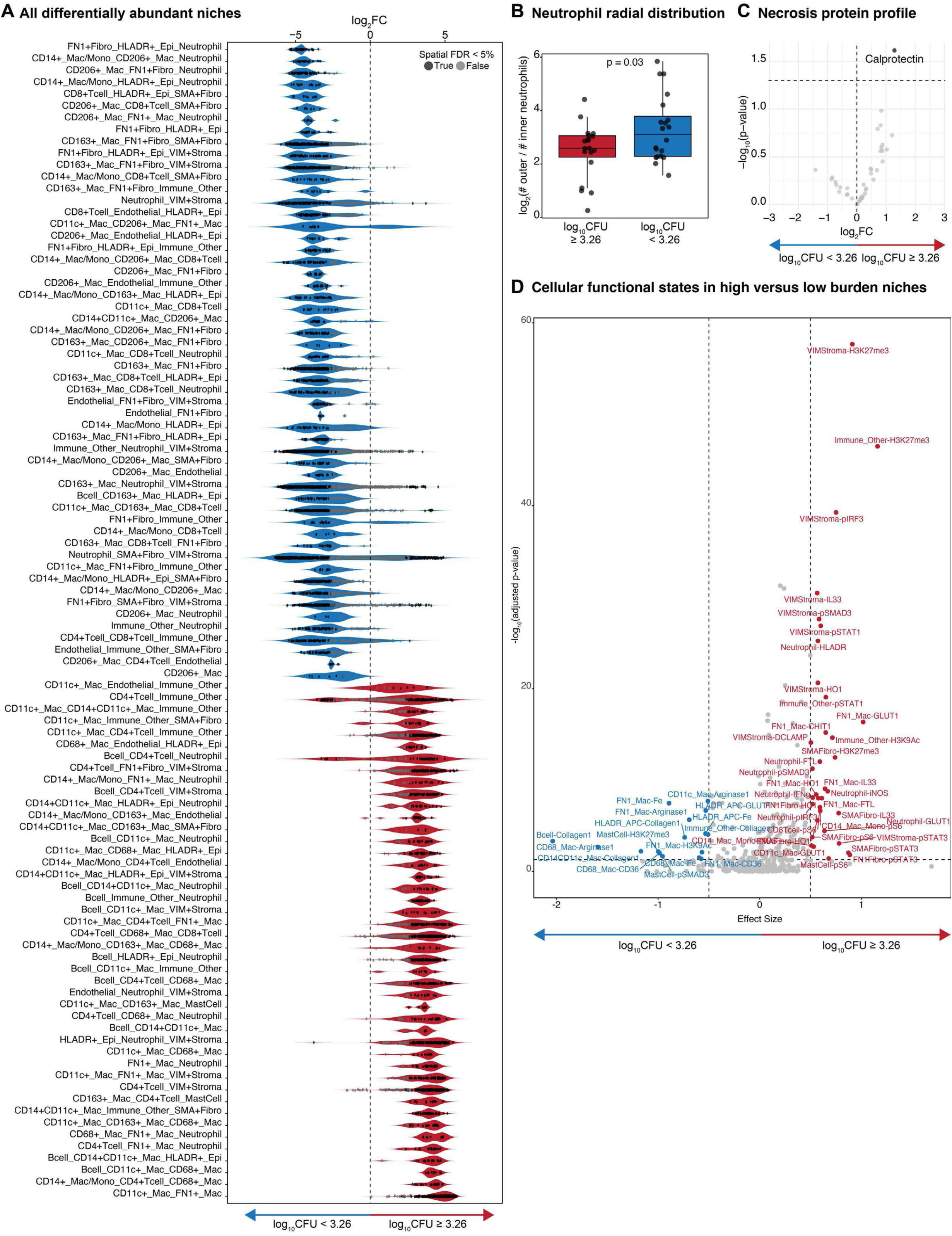
Cellular niche and spatial network analyses of TB granulomas. All significantly differentially abundant niches in low (blue) versus high (red) bacterial burden granulomas. Violins display the log fold-change with dots colored by statistical significance (determined by spatial FDR < 5%). **B)** Ratio of neutrophils in the outer versus inner region of the granuloma in low versus high bacterial burden granulomas. Boxplots display the median and IQR (25–75%), with whiskers representing the upper- and lower-quartile ±1.5× IQR. **C)** Volcano plot displaying differential expression (log_2_ fold-change) of proteins in the necrotic zones of granulomas with high or low bacterial burden. Horizontal dashed line indicates adjusted p-value < 0.05. **D)** Volcano plot displaying the effect size for all cell type-protein pairs in high (right) versus low (left) burden spatial niches. Unless specified, p-values were calculated with t tests.

**Figure S11:**
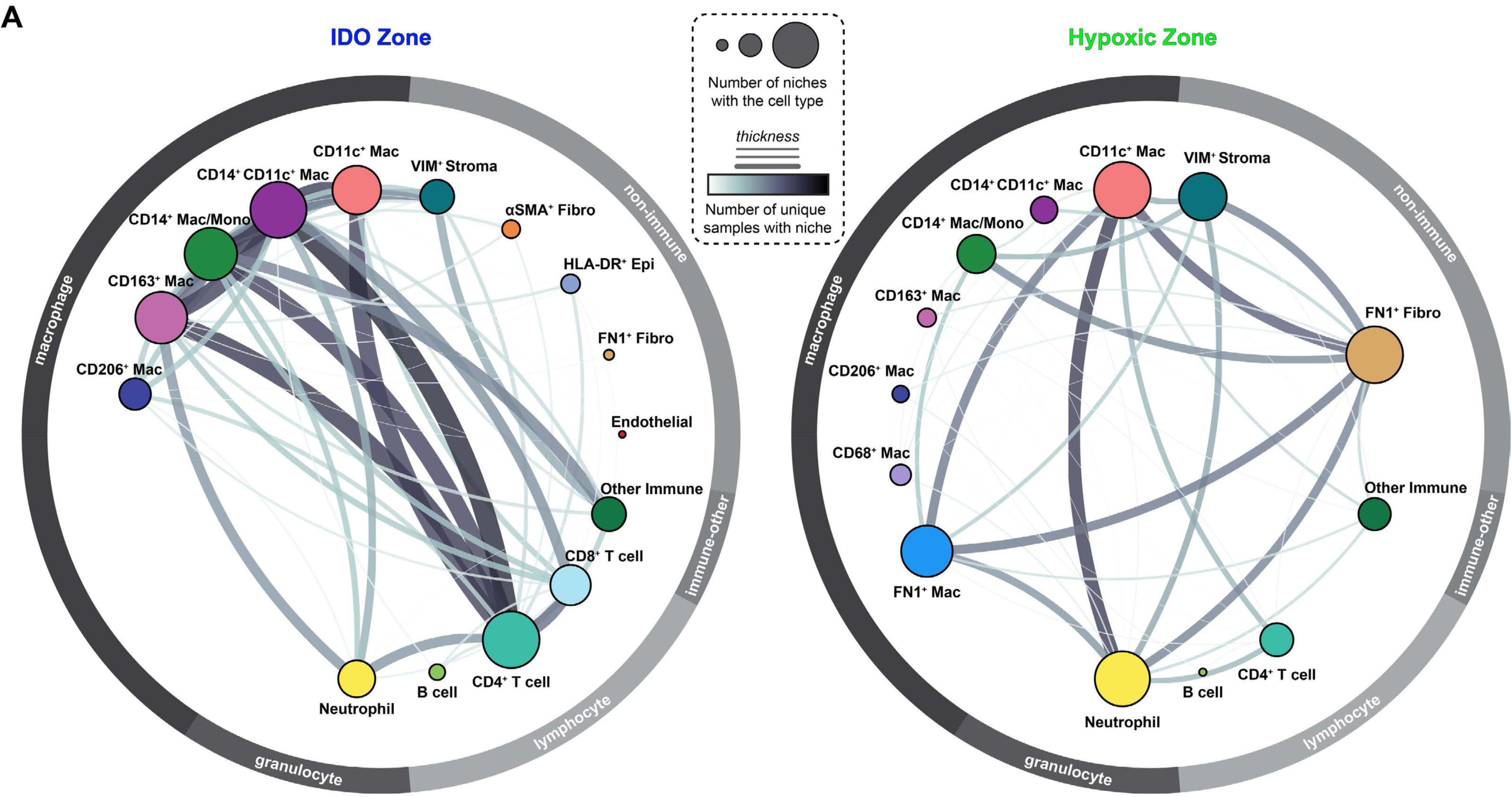
Cellular niche network of myeloid core metabolic zones. **A)** Cellular networks derived from differentially abundant niches in the IDO (left) versus Hypoxic (right) zones. Each node represents a cell type. Node size is proportional to connectedness, as measured by eigenvector centrality. Edge thickness is proportional to the number of unique samples with the corresponding interaction.

